# Decoding Musical Valence And Arousal: Exploring The Neural Correlates Of Music Evoked Emotions And The Role Of Expressivity Features

**DOI:** 10.1101/2024.02.27.582309

**Authors:** Alexandre Sayal, Ana Gabriela Guedes, Inês Almeida, Daniela Jardim Pereira, César Lima, Renato Panda, Rui Pedro Paiva, Teresa Sousa, Miguel Castelo-Branco, Inês Bernardino, Bruno Direito

## Abstract

Music can convey basic emotions, such as joy and sadness, and more complex ones, such as tenderness or nostalgia. Its effects on emotion regulation and reward have attracted much attention in cognitive and affective neuroscience. Understanding the neural correlates of music-evoked emotions may guide the development of neurorehabilitation interventions based on music. Here, we used fMRI to examine the relationship between the classification of music excerpts regarding perceived valence and arousal and their neural correlates. Twenty participants were scanned while listening to 96 musical excerpts, which were classified beforehand into four categories as a function of valence (positive vs. negative) and arousal (high vs. low). Differences in valence and arousal modulated activity in cortical regions, most noticeably the music-specific subregions of the auditory cortex, but also in the thalamus and regions of the reward network such as the amygdala. Using multivoxel pattern analysis, we created a computational model able to decode the valence and arousal of the music excerpts significantly above chance. We further explored how a set of musical features relate to brain activity in valence-, arousal-, reward-, and auditory-related ROIs. The results emphasize the differential involvement of musical features in the brain, notably expressive features such as Vibrato and Tonal and Spectral dissonance in valence, arousal, and reward brain networks, while a broader set of features modulate sensory auditory networks. Using ecologically valid music stimuli, we contribute to the definition of the neural substrates of music listening and evoked emotions. Moreover, the definition of the musical features that modulate specific brain networks paves the way to developing novel music-based neurorehabilitation strategies.

## 1 Introduction

Music can elicit a wide range of affective responses in the listener, from broader positive or arousing feelings to more specific emotions such as joy, sadness, or nostalgia. The precise mechanisms through which music induces emotions, the characteristics of these emotions, and their connection to other affective processes are the focus of much research [1]. Understanding the relationship between musical expressivity and affective responses is essential to developing effective and personalized strategies for emotion regulation [2], [3].

The effects of music on emotion regulation and reward have attracted plenty of attention among cognitive neuroscientists. Functional neuroimaging studies on music and emotion have demonstrated the involvement of emotion-related brain structures, such as the amygdala, nucleus accumbens (NAcc), hypothalamus, hippocampus, insula, cingulate cortex, and orbitofrontal cortex. In particular, a meta-analysis [4] showed that music-evoked emotions with positive valence engaged the entire reward network, consisting of the ventral and dorsal striatum (head of the caudate nucleus), amygdala, anterior cingulate cortex (ACC), orbitofrontal cortex (OFC), insula, mediodorsal thalamus (MD), and secondary somatosensory cortex (SII).

The involvement of the hippocampus in music-evoked emotions such as joy and tenderness and endocrine effects, suggests its role in emotional function, and formation and maintenance of social attachments [5]–[7]. Moreover, the hippocampal-auditory system is also key to long-term auditive memory. Its connections to the reward structures further implicate it in the emotional processing of music [8].

[9] extended the functional profile of the auditory cortex and hypothesized an emotion-specific functional hub with connections with a broad range of limbic, paralimbic, and neocortical structures.

Listening to music involves tracking and predicting sound events over time. These are key mechanisms that consistently engage the reward system [10], [11]. Both the perceptual prediction error (predictions about the music itself, i.e., the next chord) and the reward prediction error (how emotionally rewarding a piece of music is) play a role in the pleasure potential of music [12], [13].

Music proficiently exploits our expectations by manipulating melody, rhythm, and patterns [14]. The relation between specific musical features and the recognized musical emotions has been previously studied as a correspondence model based on acoustic features (such as melody, rhythm, harmony, dynamics, or timbre) and formal semantic annotations [15], [16]. To develop technological music-based therapeutic applications (e.g. using brain-computer interfaces [17], [18]), we need to identify the neural correlates of music-evoked emotions and understand how music can convey meaning to the listener [19], [20].

Different theoretical approaches to emotion and music-evoked emotion have been proposed [21], with relevance to emotion generation and regulation [22]. Valence and arousal (core affect [23]) can be taken as consciously accessible emergent properties of more basic physiological processes and neural correlates, reflecting low dimensional descriptive features of all emotional experiences [24].

The present study addresses the relation between the classification of music excerpts regarding perceived affective states and the associated neural correlates, as measured by functional magnetic resonance imaging (fMRI). Using multivariate pattern analysis (MVPA), we aim to address the relation between music characteristics and valence and arousal, identifying the brain regions that encode this information.

Previous studies using fMRI were able to successfully decode music-based emotions and explored different elements of the stimuli, such as the duration of music excerpts, decoding accuracy, and the spatial distribution of important voxels. Encoding models based on the temporal and frequency dimensions of musical stimuli (musical features encompassing rhythmic, timbral and tonal properties) revealed a spatial and temporal structure of the underlying neural representations, and decoding performance accuracies were as high as 95% [25]. Neocortical clusters in the ACC, insula and somatosensory cortex were key in the decoding of emotions in music stimuli, particularly joy and fear [26]. Decoding models were also used to identify genres of songs (described by the features ‘melody schema’ (absolute pitch, relative pitch), and ‘acoustic chromagram’ (absolute pitch)) considering voxel pattern responses [27].

Here, we aim to identify the brain networks that respond to music-evoked emotions, specifically to low-level dimensions of emotion (valence and arousal), and to understand how acoustical features of the stimuli impact their activity. Ultimately, we aim to establish the connection between music descriptors, music-derived affective states, and brain activity patterns, using machine learning applied to fMRI. We hypothesize that the neural networks involved in affective processes [28], which include emotion-dependent frontal regions, supplementary motor areas, and subcortical regions such as the amygdala, hippocampus, and striatum, in coordination with the auditory cortex, contribute to the mapping of music features and evoked emotions. Many studies have focused on very restricted music stimuli (few genres and/or music excerpts) and usually use mass-univariate approaches. Here, we propose using a more comprehensive and representative dataset, which includes several musical genres. These more naturalistic stimuli, combined with the exploratory framework proposed, will allow us to explore the relation between low-level emotional components and patterns of brain activity.

## 2 Methods & Materials

### 2.1 Participants

Twenty individuals (11 females; age range 21-41 years, M = 31.3, SD = 6.5) participated in the experiment. All participants gave written informed consent. The study was conducted following the declaration of Helsinki and approved by the Comissão de Ética e Deontologia da Investigação da Faculdade de Psicologia e Ciências de Educação da Universidade de Coimbra. Exclusion criteria included diagnoses of a neurological or mood disorder and hearing disorders.

The mood states of the participants were characterized using the Profile of Mood States (POMS) questionnaire [29], which includes 42 words describing sensations that people feel in everyday life. Participants were asked to select an answer based on what best corresponds to the way they have been feeling - a scale from 0 (Not at all) to 4 (Very much). Mood state was analyzed according to six subscales - Tension (Mdn= 7.5, range: 2-19), Depression (Mdn= 1.5, range: 0-15), Hostility (Mdn= 2, range: 2-15), Vigor (Mdn= 13.5, range: 5-22), Fatigue (Mdn= 5.5, range: 0-15), and Confusion (Mdn= 5, range: 1-17). A Total Mood Disturbance score revealed a Mdn= 109.5 with a range from 88 to 162. Our sample reported 2.2 ± 3.5 years of formal musical training and scored 18.1 ± 4.6 in the Mini-PROMS (Profile of Music Perception Skills) test [30], [31].

### 2.2 Music dataset and acoustical feature analysis

The stimuli were selected from a public dataset - 4Q audio emotion dataset [16] - of 900 30-second audio clips, divided into four categories or quadrants: Q1 (positive valence and high arousal); Q2 (negative valence and high arousal); Q3 (negative valence and low arousal); and Q4 (positive valence and low arousal). The dataset was built using a semi-automatic selection and classification method and includes 225 music excerpts per category. The dataset also includes 1702 features for each clip, both acoustic (such as tempo change and loudness) and musical (such as the number of musical layers, tremolo notes, or vibrato rates). The authors used available audio toolboxes to generate features and created novel, emotionally-relevant audio features based on previous studies. Using statistical classifiers, [16] could discriminate valence and arousal based on these features and extract relevant information regarding music emotion recognition, namely the weight of specific features and musical concepts to each emotion quadrant.

### 2.3 Procedure

Before scanning, participants filled out the MRI safety questionnaire that verified the participants’ compliance with the MRI environment, and the POMS questionnaire. The fMRI session had a total duration of approximately 75 minutes and was divided into four runs. Participants were asked to perform the listening task with their eyes closed. Each participant listened to 96 musical excerpts (24 per run) randomly selected from the dataset and balanced across the four quadrants. The excerpts were reduced in duration (11.5 seconds) by selecting the first 11.5 seconds available. Within each run, participants listened to 24 music clips, 6 of each quadrant, grouped into two sets of 3 stimuli (each block lasted 36 seconds - 11.5 seconds per excerpt and 0.5 seconds intervals in between). Between the music blocks, participants listened to 12 seconds of ambient sound (inherent to an MRI scanner), then 12 seconds of white noise, and again 12 seconds of ambient sound (Figure 1). The run structure was based on the randomized presentation of music from the four quadrants (two trials) interleaved with blocks of ambient sound (no auditory stimulus) and of white noise. Within each block, all stimuli were presented in pseudo-randomized order so that the second pass of music from a quadrant occurred only after the first presentation of all quadrants. The total duration of each run was 10 minutes.

**Figure 1.**
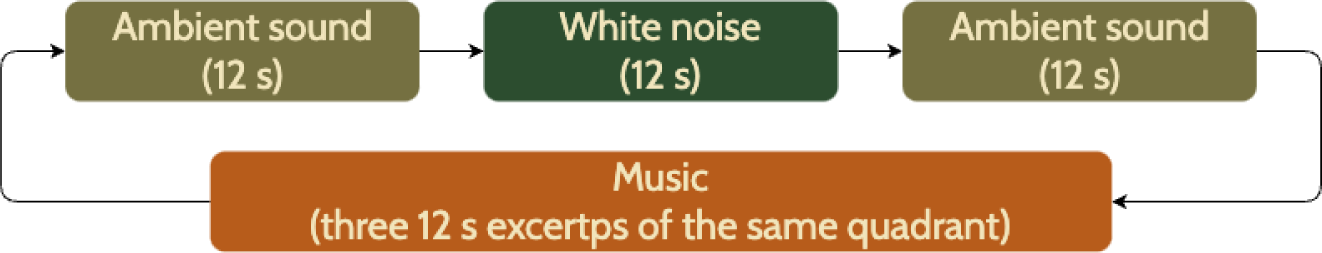
Diagram of the fMRI paradigm trial. Each trial was repeated twice for each of the four quadrants.

After the scanning session, each participant performed a rating procedure: they were asked to rate their perceived valence and arousal while listening to the music excerpts that were previously presented during the MRI session (the presentation of the music excerpts was randomized per run). The instruction specifically asked the participants to classify the stimulus in terms of both arousal and valence bipolar dimensions: “While listening to the music, you should evaluate what the music conveys to you. As soon as you think you can classify the musical excerpt in terms of valence and activation, you should click with the mouse on the point in the space, minding that the relative position to the center should also be considered”. The answer was registered as a mouse click in the 2D arousal-valence plane (Figure 2) and the click should be made when the participant identified the position. A 1-second interval followed each click, with the next music excerpt being then presented. The task was performed with noise-canceling headphones in a quiet room.

**Figure 2.**
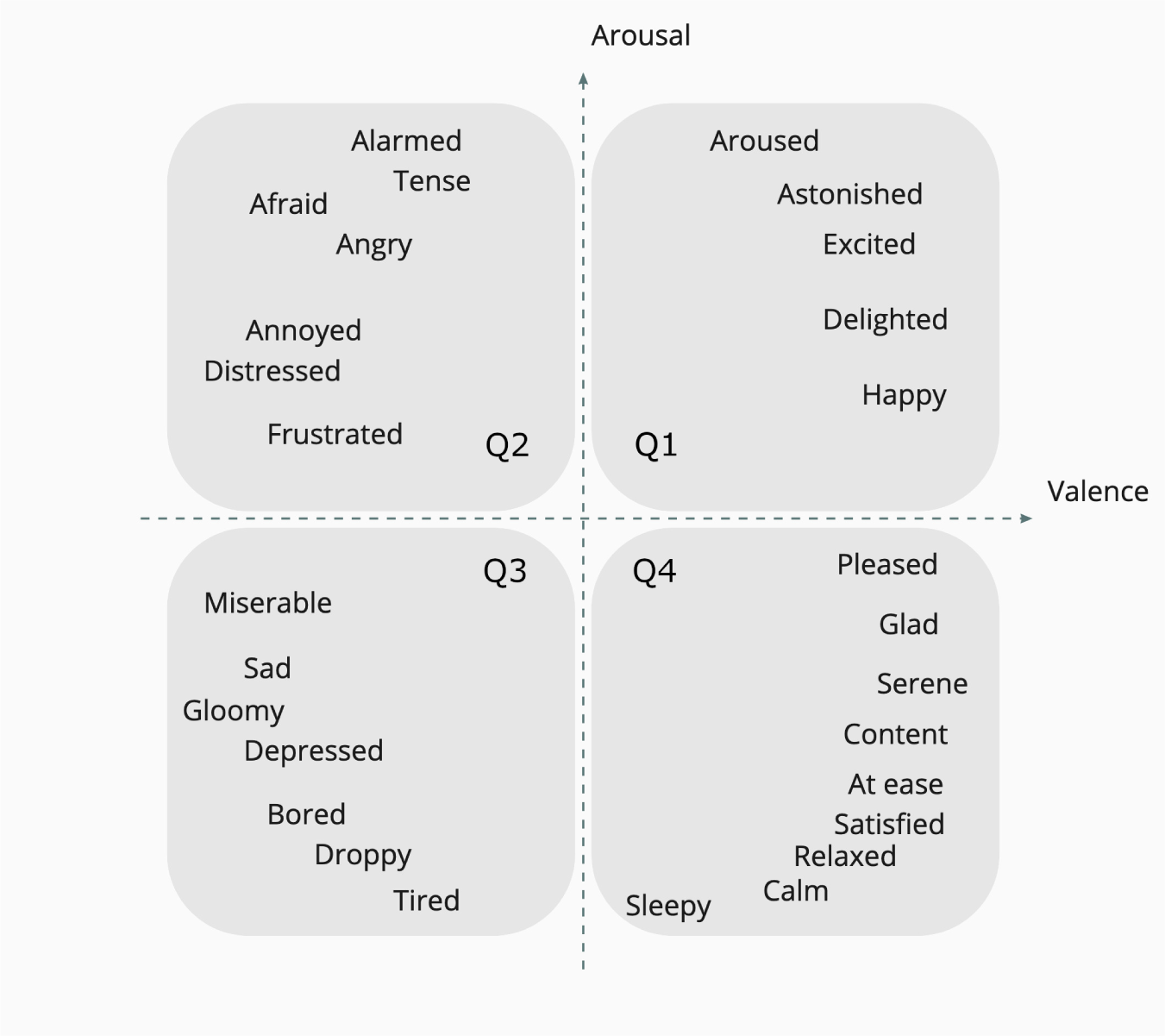
Image of the arousal-valence plane shown to the participants during the rating experiment (the text with the example emotions disappeared after experiment start). Q1 (positive valence, high arousal), Q2 (negative valence, high arousal), Q3 (negative valence, low arousal), and Q4 (positive valence, low arousal).

### 2.4 Data acquisition

The MRI session and the rating experiment were carried out using Psychopy version 2022.2.5 [32]. POMS responses were provided by adapting the 42-item paper questionnaire to a digital form.

MR acquisition was performed with a 3 T Siemens Magnetom Prisma fit scanner with a 20-channel head coil at the Institute of Nuclear Sciences Applied to Health, Coimbra. Auditory stimuli were presented using MRI-compatible headphones (Optoacoustics Optoactive II). The scanning session started with the acquisition of two anatomical images: one 3D anatomical magnetization-prepared rapid acquisition gradient echo pulse sequence (repetition time (TR) = 2530 ms, echo time (TE) = 3.5 ms, flip angle (FA) = 7°, 192 slices, voxel size 1.0 × 1.0 × 1.0 mm, field of view (FOV) = 256 × 256 mm) and one T2 space sequence (TR = 3200 ms, TE = 408 ms, 192 slices, voxel size 1.0 × 1.0 × 1.0 mm, FOV = 250 × 250 mm). A total of four functional runs were acquired using a 2D multi-band (MB) gradient-echo echo-planar imaging sequence (TR = 1000 ms, TE = 37 ms, flip angle = 68°, 600 volumes, 66 slices, voxel size 2.0 × 2.0 × 2.0 mm, FOV = 220 mm, MB factor = 6). For correcting image distortions related to magnetic field inhomogeneities, we acquired pairs of spinecho images with anterior-posterior and posterior-anterior phase encoding polarity with matching geometry and echo-spacing to each of the functional scans (TR = 10250 ms, TE = 73.6 ms). These were acquired before the functional runs. The neuroimaging data were organized according to the BIDS specification [33], using BIDSkit [34] and dcm2niix [35] tools for conversion.

### 2.5 Data analysis

Ratings of valence and arousal for each of the 24 music excerpts ranged from −1 to 1, given they were presented together in the same plane. We estimated the distribution of the ratings provided by each participant (2D position in the valence-arousal plane) for the music excerpts previously heard during the fMRI session. We report the match between the affective labels and the participants’ evaluation.

Acquired imaging data were preprocessed using fMRIPrep 20.2.7 [36], [37], which is based on Nipype 1.7.0 [38], [39]. In brief, this pipeline includes slice time correction, head motion correction, unwarping, and normalization to the MNI space. For a step-by-step description of the methods, see supplementary materials. The following sections describe the imaging data analyses performed in Python with Nilearn v0.10.2 [40].

#### 2.5.1 GLM analysis

The first level analysis was based on a single general linear model (GLM) estimated for each participant. The design matrixes included predictors for the music of each quadrant according to the participant’s labeling and confound predictors for head motion (six motion parameters + first-order derivatives + powers, a total of 24). Before generating the maps, we applied temporal high pass filtering with a cut-off frequency of 0.007 Hz, spatial smoothing with a Gaussian kernel of full-width at half maximum = 4 mm, and considered a second-order autoregressive model AR(2) as the temporal variance model.

The second level analysis consisted of generating the group activation map for the contrast between all the quadrants and white noise, corrected for multiple comparisons at the voxel level using false discovery rate (FDR) (q=0.005). We extract the clusters from this map considering a minimum cluster size of 25 voxels. Based on these clusters, we extracted its z-values for the contrast between each quadrant and noise for each participant. We only considered the voxels inside each cluster mask that were significant at the subject level for the contrast between all quadrants and noise (FDR q=0.05, k>10). We compared the z-values between each pair of quadrants for all clusters using a two-sided Mann-Whitney-Wilcoxon test with FDR correction (q=0.05). All clusters were labeled according to two atlases - AAL3 and Neurosynth - and manually confirmed by a neuroradiologist.

#### 2.5.2 Multivoxel decoding analysis

For each subject, a single GLM was re-performed on the preprocessed functional image, in which each trial of each condition of interest is separated into its own condition within the design matrix. As our conditions of interest are the music excerpts, we defined a predictor for each music excerpt labeled according to the class, i.e. the quadrant in Russell’s circumplex, identified by the participant in the rating procedure [41].

The model results in 96 beta maps (24 music excerpts per run, four runs). The individual beta maps were masked using a gray-matter (GM) mask obtained from the fMRIPrep preprocessing of the anatomical images, normalized to zero mean, and scaled to unit variance. Then, a within-subject analysis was performed on these beta maps.

Multivariate pattern analysis was carried out using a supervised learning method as the estimator model - a Support Vector Machine (SVM) with a linear kernel and an L1 regularization factor. To solve the multiclass problem, we implemented a one vs. all method.

We implemented cross-validation to split the data into different sets, we can then fit the estimator on the training data set and measure an unbiased error on the testing set. To avoid any temporal leakage between training and testing samples (bias due to temporal proximity of samples, particularly relevant in functional data), we implemented a leave-one-run-out (LORO) strategy, where the classifier was trained on 3 of the runs and tested on the remaining one. Leaving out blocks of correlated observations, rather than individual observations, is crucial for non-biased estimates [42].

##### Prediction accuracy at chance

When performing decoding, the prediction performance of a model can be checked against null distributions or random predictions. For this, we guess a chance level score using simple or random strategies while predicting condition *y* with *X* imaging data. In this work, we used dummy predictors implemented in Nilearn, to estimate a chance level score. In summary, the *DummayClassifier* function makes a prediction that ignores the input features by respecting the class distribution of the training data. This allows us to compare (two-sample t-test) whether the model is better than chance or not while still considering calls imbalance.

##### Visualizing the decoder’s weights

With linear models often used as decoders in neuroimaging, model averaging is appealing as it boils down to averaging weight maps. The weight map spatially represents the model’s weights by showing the contribution of each voxel in the image. The weight maps for the SVM model were obtained for each participant, where we highlighted the top 2% of the sorted distribution of the weights. We used AAL3 to identify the brain structures where clusters were identified [43].

#### 2.5.3 Exploring the importance of acoustic features using regression analysis

Previous research using controlled paradigms played an important role in investigating the neural processing of individual musical features and generating hypotheses for further studies [44]. In this work, we aim to comprehensively explore the relationship between acoustic features (particularly their emotional relevance) and neural correlates. Two factors contribute to this challenge - the number and diversity of the music excerpts used and the set of acoustic features. Here, we considered the top 100 features defined by [16] as the optimal set to discriminate valence and arousal.

To appropriately define the relevance of each feature in the valence, arousal, auditory processing, and reward-identified networks, we defined brain masks based on i. large-scale meta-analysis data provided by Neurosynth [45], using the “uniformity-based test” feature (the terms used for the four masks were valence, arousal, primary auditory), and ii. cluster 1 from the meta-analysis presented in [4] as a music-specific reward mask. We intersect these brain masks with the GM mask. To compute the regression coefficients between the beta series and the value of the acoustic features, we used *DecoderRegressor* method implemented in Nilearn with ridge regressor as the estimator. It implements a model selection scheme that averages the best models within a cross-validation loop (set as 3 folds). The resulting average model is the one used as a regressor. To assess the statistical significance of the results, we performed an exploratory one-sample Wilcoxon test using the coefficients obtained (uncorrected).

## 3 Results

The following paragraphs present the four sets of findings based on the analysis plan presented above. Firstly, we briefly describe the rating task and compare its results with the quadrant labeling of the music dataset. Secondly, we describe the neural correlates of music listening following a GLM approach, and present z-values per condition to characterize valence and arousal in the regions of interest (ROIs). Thirdly, we show the machine learning approach taken to differentiate the four quadrants and present the spatial distribution of the model weights, i.e., the most important voxels for decoding. Finally, we explore the functional patterns of valence, arousal, primary auditory, and reward networks as probed by acoustic features.

### 3.1 Music excerpt rating in 2D valence and arousal plane

Behavioral data results of the rating experiment are summarized in Figure 3. The results show some level of mismatch between the labels attributed by the participants and the labels available in the 4Q audio emotion dataset. This is particularly relevant in the low arousal labels (Q3 and Q4), in accordance with the dataset’s authors [16].

**Figure 3.**
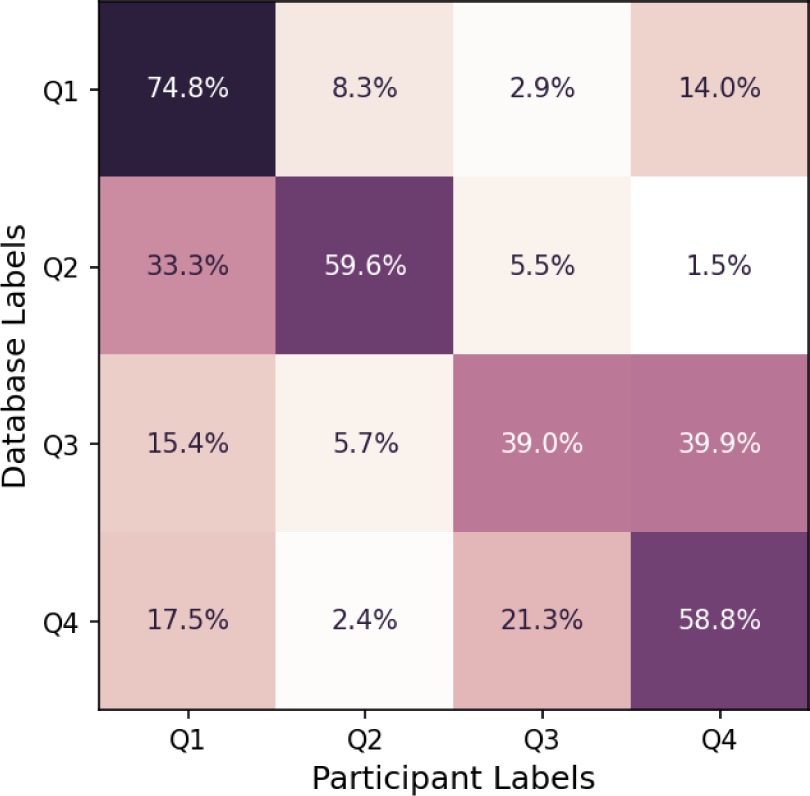
Confusion matrix comparing the labels originally associated with the music clips in the database with the labels posteriorly attributed by the participants to each music clip. The value in each cell indicates the percentage of participant ratings over the total number of music excerpts of each quadrant according to the dataset labels.

The participants’ labels for Q1 show that 74.8% of the ratings match the 4Q audio emotion dataset. A lower agreement was detected for Q3 music excerpts (low arousal and negative valence), where only 39.0% of the ratings matched with the 4Q audio emotion dataset, consistent with participant-specific interpretations of the emotional content of the songs. Based on the existence of these disparities, we performed all the following analyses using the participants’ labels.

Figure 4 depicts the differences between prior labeling and participants’ ratings for arousal and valence. The results show that Q3 stimuli (considering the 4Q audio emotion dataset labeling) were perceived as neutral here. On the opposite, the arousal overall pattern generally follows the prior labeling.

**Figure 4.**
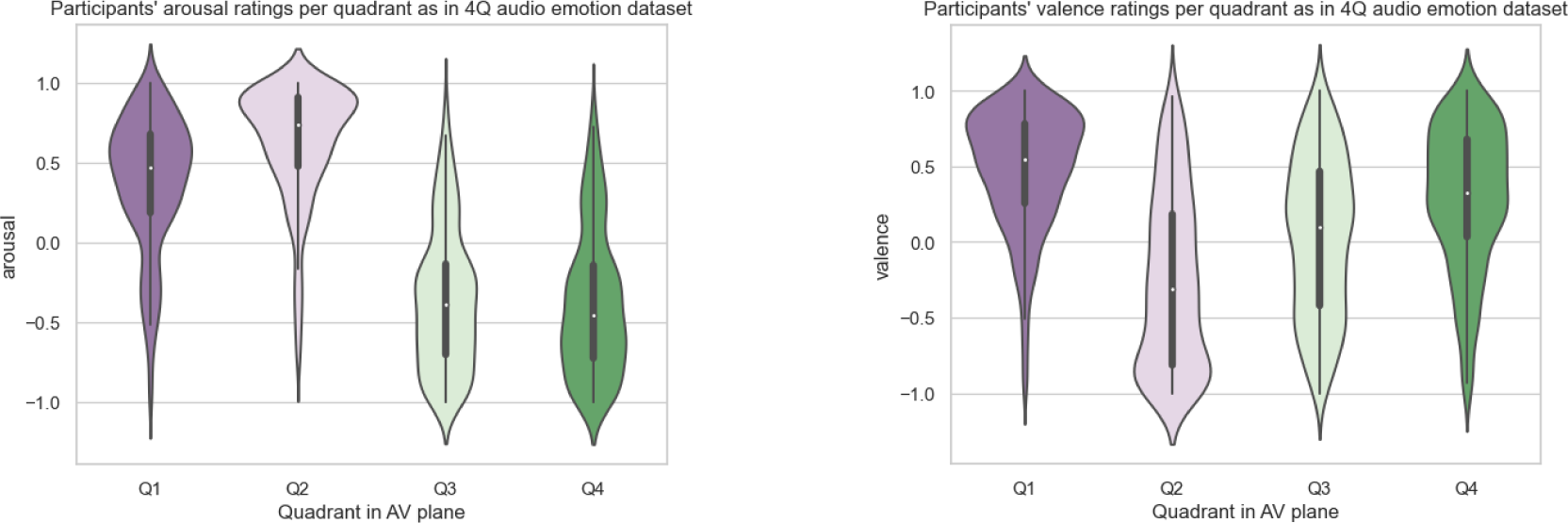
Violin plots displaying the symmetric kernel density estimate and the quartiles of a box plot of the participants’ rating for the music of all quadrants in the arousal-valence plane. The values displayed are the Euclidian distance from each axis to the position in the plane where the participants clicked normalized for the furthest report.

### 3.2 Neural correlates of music and music-evoked emotions

In Figure 5, we show the group activation map for the contrast between listening to music of all quadrants and noise. We found clusters in the auditory cortex, thalamus, cerebellum, and motor cortices, but also regions of the reward network such as the amygdala and putamen. The full list of clusters can be found in Table 1 and the z values of each identified cluster divided per quadrant are shown in Figure 6. This figure highlights activation differences between quadrants and noise in the auditory cortices, thalamus, cerebellum, and SMA, with Q1 music eliciting more activity in the majority of the clusters. Additionally, we provide the z-scores contrasting positive and negative arousal and valence in supplementary Figures S14 and S15, respectively.

**Figure 5.**
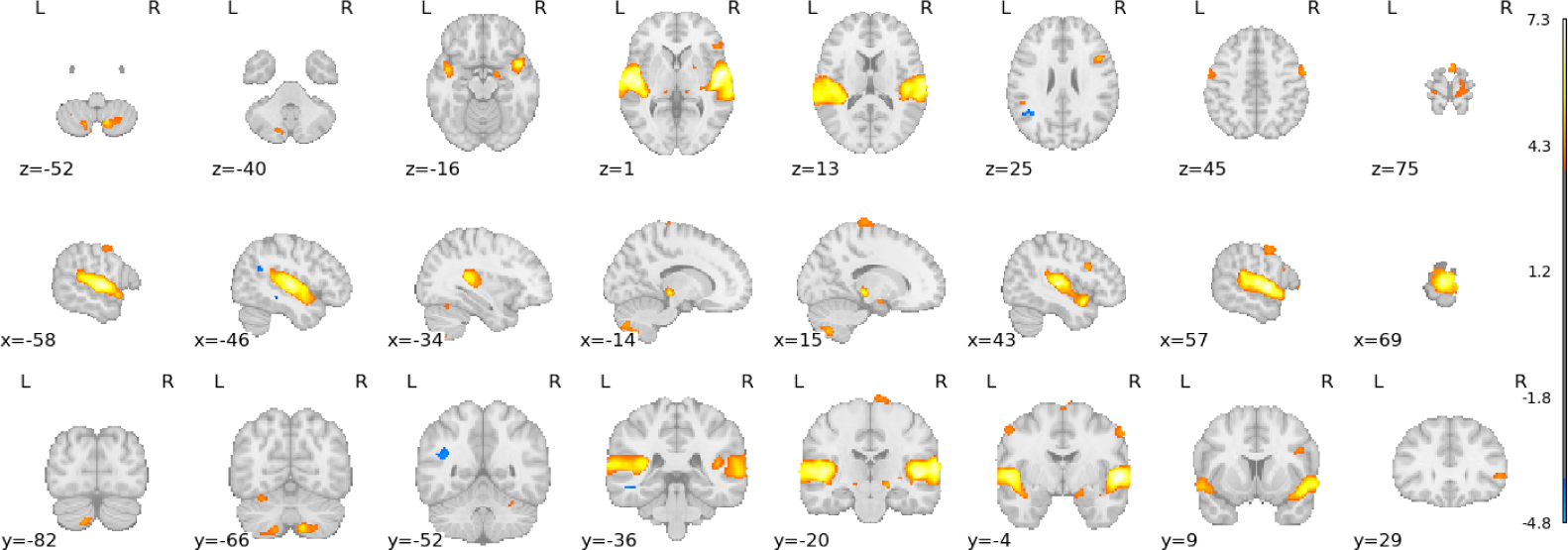
Group activation map for the contrast between all quadrants and noise (z-values FDR corrected, q = 0.005, cluster correction k > 25, N=19).

**Figure 6.**
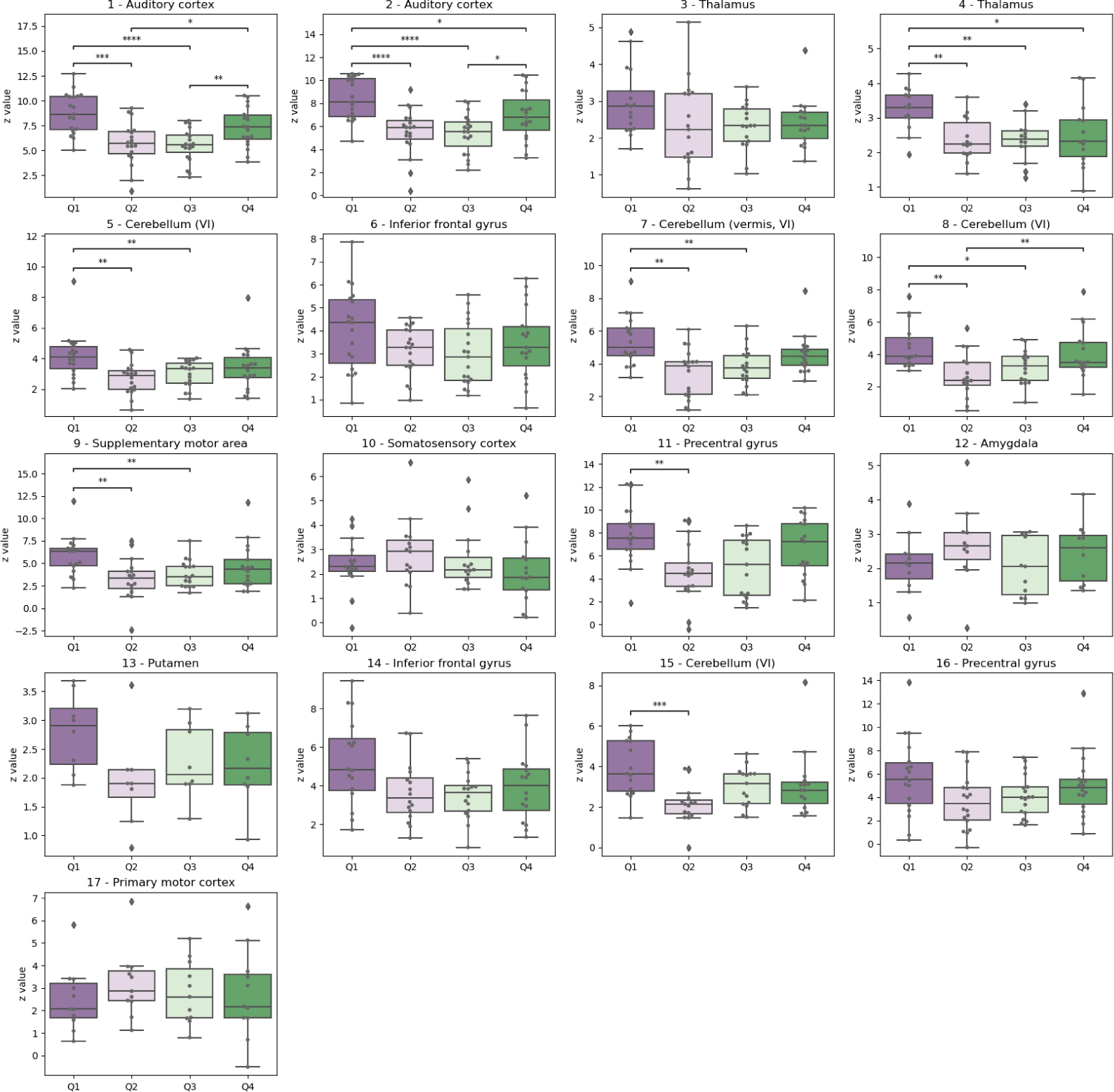
Boxplots for the z values of each identified cluster (Table 1) divided per quadrant. Pairwise statistical comparisons between quadrants are displayed, performed using two-sided Mann-Whitney-Wilcoxon tests with FDR correction (q=0.05).

**Table 1.** Cluster table of the group activation map contrasting music from all quadrants and noise (FDR corrected, q = 0.005, cluster correction k > 25, N=19). The labels for major clusters were obtained from the AAL3 atlas and the Neurosynth association tab.

| Cluster ID | X | Y | Z | Peak z value | Cluster Size (mm3) | Labels |
| --- | --- | --- | --- | --- | --- | --- |
| 1 | 58 | -5 | 2 | 7.29 | 34968 | Temporal superior sulcus, auditory cortex |
| 1a | 50 | -15 | 4 | 7.05 |  |  |
| 1b | 66 | -29 | 8 | 6.89 |  |  |
| 1c | 44 | -25 | 6 | 6.82 |  |  |
| 2 | -59 | -11 | 4 | 7.01 | 34376 | Temporal superior sulcus, auditory cortex |
| 2a | -49 | -17 | 2 | 6.95 |  |  |
| 2b | -45 | -23 | 8 | 6.75 |  |  |
| 2c | -51 | -1 | -3 | 6.43 |  |  |
| 3 | 16 | -25 | -5 | 5.84 | 952 | Thalamus |
| 4 | -15 | -27 | -7 | 5.50 | 816 | Thalamus |
| 5 | 12 | -67 | -53 | 5.35 | 2376 | Cerebellum (VI) |
| 5a | 30 | -61 | -47 | 4.89 |  |  |
| 5b | 12 | -77 | -47 | 4.56 |  |  |
| 6 | 48 | 16 | 22 | 4.97 | 1040 | Inferior frontal gyrus |
| 6a | 60 | 16 | 24 | 3.81 |  |  |
| 7 | -13 | -81 | -45 | 4.65 | 2416 | Cerebellum (vermis, VI) |
| 7a | -15 | -69 | -53 | 4.54 |  |  |
| 7b | -29 | -61 | -57 | 4.30 |  |  |
| 7c | -15 | -75 | -51 | 4.20 |  |  |
| 8 | -29 | -61 | -25 | 4.57 | 984 | Cerebellum (VI) |
| 9 | 2 | 2 | 76 | 4.56 | 1272 | Supplementary motor area |
| 10 | 14 | -25 | 82 | 4.50 | 2104 | Somatosensory cortex |
| 11 | 60 | -5 | 46 | 4.40 | 1096 | Precentral gyrus (face, mouth) |
| 12 | 18 | -7 | -15 | 4.40 | 360 | Amygdala |
| 13 | 24 | 2 | 6 | 4.37 | 312 | Putamen |
| 14 | 52 | 30 | 2 | 4.32 | 520 | Inferior frontal gyrus |
| 15 | 30 | -57 | -25 | 4.23 | 408 | Cerebellum (VI) |
| 16 | -57 | -5 | 48 | 4.18 | 800 | Precentral gyrus (face, mouth) |
| 17 | -17 | -29 | 78 | 3.95 | 264 | Primary motor cortex |

### 3.3 Music quadrant identification from its fMRI signature

In the following classification analyses, we considered data from 15 participants. As the classes were defined according to each participant’s rating of the music clips, and 4 participants did not report one of the four quadrants, their data was not included.

#### 3.3.1 Decoding results

The results for the four-class MVPA are shown in Figure 7. The mean overall accuracy was 69.4% (Q1 = 58.1%, Q2 = 75.3%, Q3 = 74.9%, Q4 = 69.3%) and the difference between the overall accuracy and the accuracy of the stringent dummy predictor is statistically significant (p<0.001).

**Figure 7.**
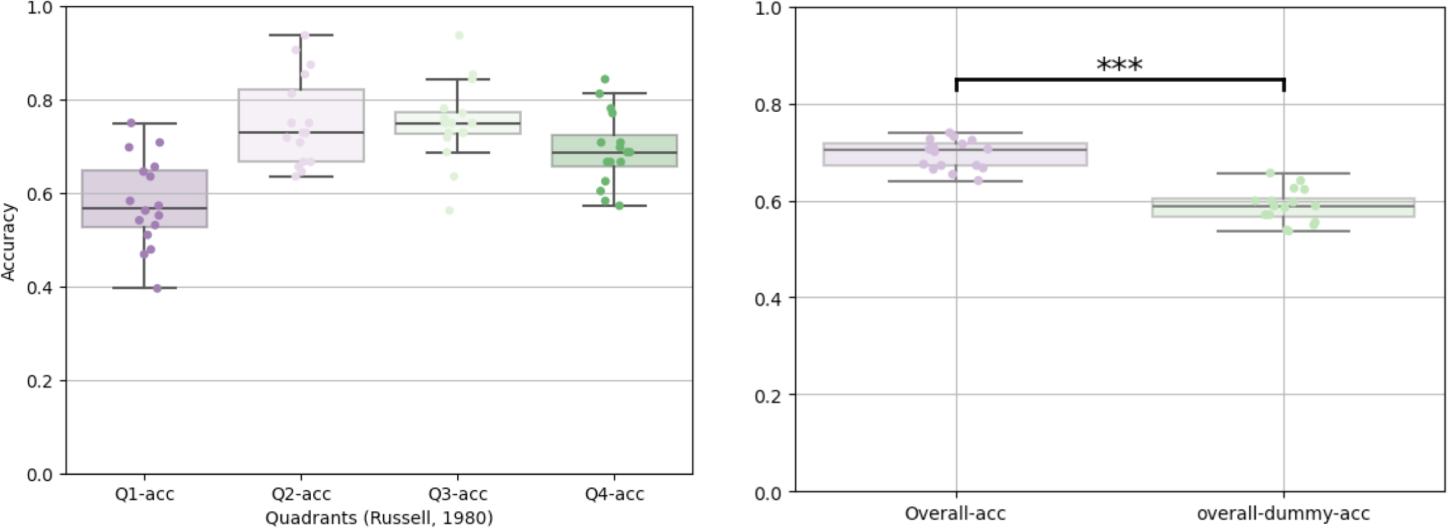
MVPA classification accuracy results. On the left, we display the boxplot of the accuracies achieved per quadrant. On the right, we compare the mean overall accuracy with the accuracy of the dummy predictors (*** - p < 0.001).

#### 3.3.2 Visualization of weights

To explore the spatial distribution of the voxels that better differentiate quadrants, we selected the top 2% most discriminant voxels for each one vs. all models (Q1 vs. other, Q2 vs. other, etc.) and overlapped them. Figure 8 presents a probability map over the four maps (cluster threshold with 50 voxels). The results show that the most important voxels were predominantly located in the bilateral auditory cortices (Figure 8) for all quadrants. The full list of clusters can be found in Table 2.

**Figure 8.**
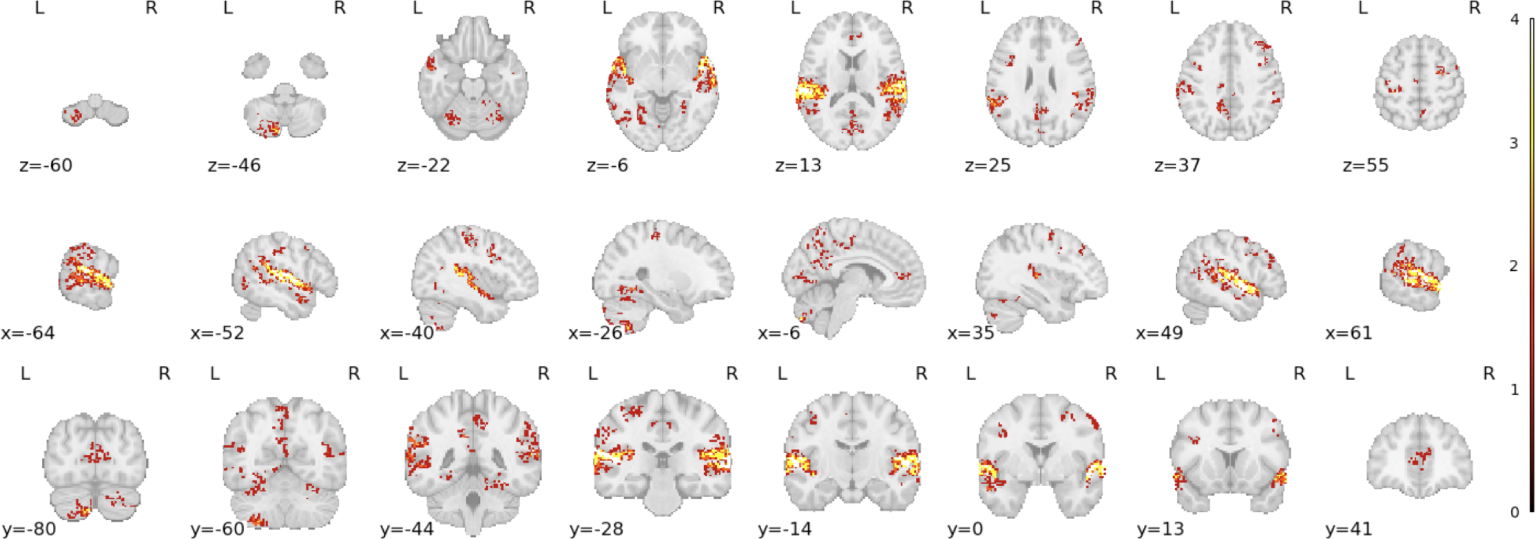
The combination of the top 2% most important voxels for each quadrant to classify valence and arousal categories in the whole-brain analysis (the colorbar represents the number of times that a specific voxel appears in the top 2% - the most important voxels for the classification task).

**Table 2.** Description of the clusters identified in the map of Figure 8, displaying the coordinates of the peak voxel, the number of quadrants at that coordinate, and the labels from the AAL3 atlas and Neurosynth associations.

| X | Y | Z | #Q at peak | Cluster Size (mm3) | Labels |
| --- | --- | --- | --- | --- | --- |
| -65 | -3 | -3 | 4 | 31904 | Temporal superior sulcus, auditory cortex |
| 56 | -41 | 12 | 4 | 29896 | Temporal superior sulcus, auditory cortex |
| -49 | -67 | -9 | 2 | 1936 | Temporal inferior, visual ventral stream |
| -47 | 8 | 38 | 2 | 1760 | Precentral gyrus, language |
| -33 | -31 | 56 | 2 | 1552 | Precentral gyrus, motor |
| -27 | -53 | -9 | 2 | 3032 | Lingual gyrus |
| -13 | -37 | 44 | 2 | 3888 | Middle cingulate cortex |
| -11 | -55 | 40 | 2 | 6664 | Precuneus |
| -3 | 44 | 8 | 1 | 1184 | Anterior cingulate cortex |
| 24 | -45 | -19 | 1 | 872 | Fusiform gyrus |
| 44 | 32 | 34 | 1 | 1400 | Frontal middle gyrus, pre-frontal cortex, working memory |
| 44 | 6 | 40 | 1 | 800 | Frontal middle gyrus, pre-frontal cortex |
| 56 | -1 | 46 | 1 | 816 | Precentral gyrus, motor |

### 3.4 Acoustic features and brain activation

To explore the impact of acoustic features in the valence, arousal, auditory, and reward networks, we investigated its regression profile with all acoustic features, by computing the mean over all voxels of each mask per participant.

#### 3.4.1 Musical features differential engagement in different ROIs

In this analysis, we included a set of 100 candidate acoustic features (based on the previous work and nomenclature by [16]). For a complete overview of these analyses, see Supplementary Materials.

The exploratory analysis considering the valence mask highlights Expressivity Features, as four of the seven acoustic features presenting statistically significant results are features of expressive techniques in music. In particular, these features are based on characteristics of Vibrato, an expressive technique used in vocal and instrumental music that consists of a regular pitch oscillation. The main characteristics are the amount of pitch variation (extent) and the velocity (rate) of this pitch variation. Similarly, we see a dominance of Expressivity features when considering the arousal mask. Regarding the auditory mask, we see a significant increase in the number of features associated with brain activation (54 features). Most features are related to Expressivity, Texture, and Tone Color. Considering the reward mask, we found that Loudness - Spectral and Tonal Dissonance, Pitch (Terhardt) - Chord Change Likelihood, and three features derived from Vibrato were the top-ranked features.

## 4 Discussion

Our main finding was that valence and arousal can be discriminated above chance level based on the neural representation of music listening to a comprehensive dataset with a broad range of music genres. Moreover, this classification is mainly based on bilateral auditory cortices with contributions from the precuneus, frontal, precentral, and cingulate cortices.

The GLM analysis showed that the music task activated sensory and emotion-related areas. The analysis of z-values per quadrant showed significant differences in activation of the bilateral auditory cortices, auditory thalamus, and supplementary motor area (SMA), depending on the quadrant.

The analysis of acoustic features and its differential recruitment of the valence, arousal, and auditory sensory networks highlighted expressivity features, particularly the Vibrato-related ones, as emotion-network modulators.

These findings contribute to further comprehending how music relates to emotions, defining music-specific regions of interest, and identifying specific musical features that modulate different brain networks. Ultimately, we pave the way for developing music-based neurorehabilitative strategies (such as neurofeedback): the inherent ability of music to modulate emotions and regulate mood could indicate that it is a strong candidate for the interface between the participants and their brain activity. Our previous findings support the strategic role of task complexity and feedback valence on the modulation of the network nodes involved in monitoring and feedback control, key variables in neurofeedback framework optimization [46].

### 4.1 Valence and Arousal ratings

The distinction between the emotion the listener feels and the one expressed by the music has been the subject of intense debate in the literature. Schubert defined the former as the internal locus of emotion and the latter as the external locus and discussed the relation between the two loci [2]. The wording considered in the rating task is framed in the assessment of the external locus. The short duration of the music excerpts used for the functional task (aimed to explore a wide range of genres and space of acoustical features) was a better fit to study from the point of view of perceived, but not felt, emotions. Nevertheless, [26] have previously concluded that feeling representations emerged within seconds, suggesting the use of shorter musical stimuli.

Three main theories are discussed - emotion contagion (both loci are part of the same emotional process, which explains why we tend to feel the emotion that music is conveying) [47], decoding (the listener’s decoding will not necessarily be the same and may be susceptible to ‘noise’), and dissociation (which explores the complex combination of emotion matches and non-matches, based on mixed cues, between and within loci). Notably, the emotion felt is often rated the same (particularly in the arousal factor) or lower than the corresponding emotion expressed by the stimuli [2]. The selection of a labeled public dataset aimed to appropriately balance the stimuli on the arousal-valence plane. The mismatch identified reinforces the need to tailor all the task elements individually. Several personal and stimulus elements may contribute to variability, such as physiological arousal, personality, and age, as well as musical features. Additionally, the existence of a positive emotional bias, extensively described in music literature and interpreted as a paradox of tragedy (deriving pleasure from something that is tragic), may also contribute to the shift of the ratings from negative to positive valence quadrants [48]–[50].

All the analyses presented in this study considered the labels provided by each participant at the cost of stimuli imbalance between classes. This limitation is minimized by the ability of models to adjust weights and decision hyperplanes per class.

Low arousal stimuli presented a significant mismatch with the public dataset labels. The ambiguity associated with low arousal has been previously described in music stimuli [51] and interpretation of other emotion-based stimuli, namely facial expressions [52]. Their findings suggest that low-arousal facial expressions are more ambiguous. Moreover, the authors suggest that this is related to a greater activation of an ambiguity/salience system subserved by the amygdala, cerebellum, and dorsal pons.

Key variables have been identified as possible confounds in the Music-Emotion relation, such as the wording and detail or task context [53]. The range of wordings found in the literature may benefit from standardization [15].

### 4.2 Neural correlates of music-induced valence and arousal

#### 4.2.1 GLM and activation per quadrant

The contrast of music vs. noise showed activation clusters in the superior temporal gyrus (STG), auditory thalamus, hippocampus, inferior frontal gyrus, SMA, motor and premotor cortices, amygdala, basal ganglia, inferior frontal gyrus, and the cerebellum.

The STG is known as a site of auditory association (and a site of multisensory integration) and thus necessarily plays a crucial role in music listening and music production [54]. As the contrast considers noise as the control condition, at least a component of the activations was not simply due to the sound. In this sense, in accordance with previous findings, the auditory cortex contributes not only to the sensory perception of the auditory stimuli but also to its decoding. The hemodynamic response in the bilateral auditory cortex reflects a differentiated response to valence, as pieces of music labeled from both Q1 and Q4 present stronger responses than the ones labeled with Q2 and Q3. Recent studies have shown that emotional/motivational processes profoundly influence the activity in the auditory cortex and that neurons in the auditory cortex can distinguish between the positive and negative valence of sounds [55]. One of the models proposed for this emotion-informed process in sensory cortices is based on functional connectivity studies that have shown that the reward value of music is correlated with increasing functional connectivity [4].

The auditory thalamus (medial geniculate body) is one of the structures of the auditory pathway and part of the thalamocortical system that provides the anatomical bases for tone and rhythm channels [56]. This structure is also involved in emotional reactions to acoustic stimuli and is part of a thalamocortical circuit that integrates context and content [5].

Frontal activity was found in the inferior frontal gyrus (pars triangularis and frontal operculum), particularly in the right hemisphere. [57] reported on the role of the inferior frontal gyrus in perceiving and feeling emotions during music listening, highlighting an increase in activity during the perceived emotion task. This region was also found in other studies based on MVPA, highlighting its ability to differentiate emotions (fear, happiness, sadness, tenderness, and liking in [58]; fear and joy in [26]).

Other activated clusters were found in the somatosensory and motor cortices, premotor regions within the precentral and postcentral gyri, as well as in the SMA. This aligns with previous studies reporting on the engagement of somatomotor regions and the SMA with auditory processing and acoustic cues, emotion perception, and the subjective experience of emotions [54], [59]. Notably, pleasurable music engaging the SMA demonstrates heightened activation of motor regions, including the SMA and cerebellum, compared to emotionally less evocative music (note that activity during pieces of music labeled as Q1 presents higher values) [60]. Music frequently prompts spontaneous rhythmic movements in listeners. Therefore, the sensorimotor activity observed in the present study may signify the interplay between perception and movement, even without overt physical actions.

The GLM also confirmed the contribution of the amygdala in the music-listening task. Moreover, the activation pattern in the amygdala did not show significant differences associated with valence or arousal of the music pieces. Koelsch and colleagues [61] found that brain activity in the amygdala was higher during joy than fear stimuli (after establishing that valence in joy stimuli was significantly higher than for fear). Our finding, on the contrary, accords with the view that the amygdala plays the role of a hub in an emotional coordination structure, engaged in processing both negative and positive stimuli [62].

The reward system plays a central role in the pleasure that most people derive from music. Our results suggest the engagement of this system: the ventral striatum (including the NAcc) integrates the limbic circuit and receives input from the cingulum, OFC, and mesial temporal structures. The dorsal striatum (including the putamen and the caudate nucleus) receives inputs from motor and prefrontal/associative areas and projects to the globus pallidus and substancia nigra. These circuits are highly interlinked and can both lead to dopamine release. As such, the striatum is one of the biological substrates of reward and emotion (pathway of dopaminergic neural responses). Pharmacological manipulation of dopamine causally demonstrated that dopamine mediates musical reward experience, in both positive and negative directions [63].

Several clusters were identified in the cerebellum. The cerebellum has been reported in most decoding studies and these results suggest its relation to music perception and execution [64]. Different parts of the cerebellum have been identified for different aspects of these processing tasks (emotion identification, pleasure in music-listening tasks, etc.) and their individual connections to the cerebral cortex as well as to the basal ganglia. It has been shown, for example, that cerebellar regions are involved in processing specific acoustical features [27], [65].

Altogether, our results reinforce the main structures previously identified in music perception and emotion identification studies. The heterogeneity of our music dataset highlights that this network represents a basis for processing music stimuli, and the quadrant specificity analysis reinforces the role of structures such as the bilateral auditory cortices, auditory thalamus, SMA, and cerebellum in decoding valence and arousal.

#### 4.2.2 Exploring the most important voxels for decoding

The spatial distribution of the most relevant voxels for decoding the different quadrants emphasizes the importance of the auditory cortex, motor areas, cingulate (anterior, middle) gyrus, and frontal middle gyrus for encoding valence and arousal. The results suggest a significant overlap between the main regions identified here and the GLM analysis.

### 4.3 How does music express different emotions? The neural correlates of acoustical features of music

Music emotion recognition has attracted attention from both music information retrieval and cognitive neuroscience, in a search for a better understanding of the mechanisms associated with evoked emotions based on music stimuli. Numerous features of music have been reported to be suggestive of discrete emotions [44].

Regarding the reward mask, we found that expressivity, loudness (tonal and spectral dissonance), and pitch were the most associated with the modulation of brain activity. The impact of dissonance and expressive features have been previously found to be relevant in the modulation of reward structures [6], [14], [65].

[66] studied physical voice features and concluded that the amygdala and auditory cortex decode the affective quality of the voice not only by processing the emotional content from previously processed acoustic features but also by processing the acoustic features themselves when these are relevant to the identification of the voice’s affective value.

Many of the significant features translate aspects of the vibrato. According to an early study, “a good vibrato is a pulsation of pitch, usually accompanied with synchronous pulsation of loudness and timbre, of such extent and rate as to give a pleasing flexibility, tenderness, and richness to the tone” [67]. An irrefutable characteristic of the singing voice [68], it is also extensively used in most musical instruments to address expressivity. Some even argue that vibrato emerged since it mimics the crying voice, and the manipulation of its extent, rate, or intensity may be linked to/identified as different emotions [69].

[25] extracted twenty-one acoustic features capturing timbral, rhythmical, and tonal properties and trained a model to predict brain activity patterns. While achieving 77% accuracy considering voxels within the Heschl’s gyrus, secondary auditory regions planum temporale, planum polare, and anterior and posterior STG, evidenced the relation between the modulation of these areas with musical features as roughness - a measure for sensory dissonance, root-mean-square energy - a loudness feature and sub-band flux frequencies between 200 Hz and 1600 Hz and negative loadings of flatness, i.e. a description of the smoothness of the frequency distribution. [70] explored six musical features (labeled by the authors as Fullness, Brightness, Activity, Timbral Complexity, Pulse Clarity, and Key Clarity) and their ability to predict brain responses while listening to music pieces. Areas in the superior temporal gyrus, Heschl’s gyrus, Rolandic operculum, and cerebellum contributed to the decoding of timbral features, while for the rhythmic features, the main areas contributing were the bilateral STG, right Heschl’s gyrus, and hippocampus. [27] used a spherical searchlight regression analysis to predict brain responses to melody and harmony features. Using searchlight (classification models considering localized spherical masks), statistically significant results were achieved in each of the four cerebral lobes, as well as in the parahippocampal gyrus and the cerebellum.

Our results show that expressivity features (particularly the ones based on Vibrato) tend to contribute more to the modulation of valence- and arousal-related brain structures (as defined by a Neurosynth meta-analysis). The connectivity between valence structures in the brain mask considered and the auditory cortex (the sensory input) is well-known [5], [71]. In this sense, our data suggests that musical features are directly or indirectly linked to the activation patterns of different brain areas linked to valence and arousal [72].

### 4.4 Limitations

We used a public dataset as stimuli to obtain a heterogeneous yet balanced dataset in valence and arousal. This attempt to use valence and arousal ratings from a previous study to establish a balance between quadrants was not successful, as the individual ratings of the participants did not fully agree, which may have minimized the power of our analysis. To maximize the heterogeneity and variety of music pieces in the dataset, we selected a relatively short segment of each piece. Nevertheless, recent evidence [26] has previously concluded that feeling representations emerged within seconds.

The statistical analysis of the acoustic features was not corrected for multiple comparisons. Considering the number of features analyzed and the exploratory nature of this section of our study, we should limit our interpretation to the trends shown by this analysis. Future confirmatory analyses will require out-of-sample validation or another dataset.

### 4.5 Conclusion

Our results indicate that several brain regions significantly encode the neural correlates of the peception of valence and arousal in a naturalistic set of music stimuli. Using a comprehensive dataset, we probed the valence and arousal main brain structures and mapped them. We also explored the potential of several acoustical features to modulate activity in key structures associated with valence, arousal, sensory, and reward networks.

The emotion contagion theory (that considers that both loci are part of the same emotional process) reinforces the relation between perceived and felt emotions. However, future works on aiming emotion felt (and neurorehabilitative approaches targeting emotion and mood regulation) should carefully address the optimization of the framework.

Previous studies indicate that familiar music elicits stronger emotional reactions and engagement of reward structures [60]. While this is a topic for future research, here we focused on the perception, not the feeling, of the arousal and valence of the music. Future studies, focusing on emotion felt should use longer musical segments and self-selected musical pieces to optimize emotional reaction. Moreover, the arousal-valence plane may not be the optimal emotion framework to explore emotion felt. Aesthetic emotions [73], for example, may represent a more appropriate music emotion model. The connectivity patterns involved in music emotion brain activation should also be addressed in future studies. [11] provided evidence that an increase in the reward value of music is correlated with increasing functional connectivity between the sensory cortex and reward brain structures as the ventral striatum/nucleus accumbens.

The understanding of the mechanisms between musical descriptors and specific brain structures (e.g. reward system) is key to optimizing therapeutic strategies. BCIs may take advantage of music stimuli to provide optimal rewarding feedback - neurally informed, immersive, and engaging music feedback depends on the characterization of the link between musical features and brain responses. Several authors have proposed features such as loudness and tempo of music pieces selected a priori as feedback as a regulator of volitional and non-volitional brain activity. Real-time adjustment of specific music features may represent an important alternative for emotion regulation of BCI [74]. Our results suggest that other acoustical features (for example Vibrato) may ultimately represent a better choice to regulate brain activity linked to valence and reward.

## 4.6 Acknowledgment

We thank all the participants who took part in this study. We also thank the remaining teams members of the Brainplayback project - André Granjo, Carolina Travassos, João Pereira - for the insightful contributions in the beggining of the project. A last word of gratitude to the team of the MRI Unit of ICNAS - Sónia Afonso, Tânia Lopes - for their support and assistance during the data acquisition.

## 4.7 Author contributions

Conceived and designed the analysis: AS, RP, RPP, CL, IA, IB, BD; Collected data: AS, BD; Contributed data or analysis tools: RP, RPP, AS, BD, AGG; Performed the analysis: AGG, AS, BD; Wrote the draft: BD, AS, AGG; Contributed to the manuscript: RP, RPP, CL, IA, IB, DJP, AGG,MCB.

## Data availability

All scripts used for the analyses mentioned in this work can be found here. The anonymized and defaced dataset can be found in BIDS format here. The data management plan regarding this project can be found at Zenodo.

