## Supplementary Materials for "Decoding Musical Valence And Arousal: Exploring The Neural Correlates Of Music Evoked Emotions And The Role Of Expressivity Features"

### 7. Supplementary materials

#### 7.1. fMRIPrep full method description

Anatomical data preprocessing

A total of 1 T1-weighted (T1w) images were found within the input BIDS dataset.The T1-weighted (T1w) image was corrected for intensity non-uniformity (INU) with N4BiasFieldCorrection (Tustison et al. 2010), distributed with ANTs 2.3.3 (Avants et al. 2008, RRID:SCR_004757), and used as T1w-reference throughout the workflow. The T1w-reference was then skull-stripped with a Nipype implementation of the antsBrainExtraction.sh workflow (from ANTs), using OASIS30ANTs as target template. Brain tissue segmentation of cerebrospinal fluid (CSF), white-matter (WM) and gray-matter (GM) was performed on the brain-extracted T1w using fast (FSL 5.0.9, RRID:SCR_002823, Zhang, Brady, and Smith 2001). Brain surfaces were reconstructed using recon-all (FreeSurfer 6.0.1, RRID:SCR_001847, Dale, Fischl, and Sereno 1999), and the brain mask estimated previously was refined with a custom variation of the method to reconcile ANTs-derived and FreeSurfer-derived segmentations of the cortical gray-matter of Mindboggle (RRID:SCR_002438, Klein et al. 2017). Volume-based spatial normalization to one standard space (MNI152NLin2009cAsym) was performed through nonlinear registration with antsRegistration (ANTs 2.3.3), using brain-extracted versions of both T1w reference and the T1w template. The following template was selected for spatial normalization: ICBM 152 Nonlinear Asymmetrical template version 2009c [Fonov et al. (2009), RRID:SCR_008796; TemplateFlow ID: MNI152NLin2009cAsym],

Functional data preprocessing

For each of the 4 BOLD runs found per subject (across all tasks and sessions), the following preprocessing was performed. First, a reference volume and its skull-stripped version were generated using a custom methodology of fMRIPrep. A B0-nonuniformity map (or fieldmap) was estimated based on two (or more) echo-planar imaging (EPI) references with opposing phase-encoding directions, with 3dQwarp Cox and Hyde (1997) (AFNI 20160207). Based on the estimated susceptibility distortion, a corrected EPI (echo-planar imaging) reference was calculated for a more accurate co-registration with the anatomical reference. The BOLD reference was then co-registered to the T1w reference using bbregister (FreeSurfer) which implements boundary-based registration (Greve and Fischl 2009). Co-registration was configured with six degrees of freedom. Head-motion parameters with respect to the BOLD reference (transformation matrices, and six corresponding rotation and translation parameters) are estimated before any spatiotemporal filtering using mcflirt (FSL 5.0.9, Jenkinson et al. 2002). BOLD runs were slice-time corrected to 0.445s (0.5 of slice acquisition range 0s-0.89s) using 3dTshift from AFNI 20160207 (Cox and Hyde 1997, RRID:SCR_005927). The BOLD time-series (including slice-timing correction when applied) were resampled onto their original, native space by applying a single, composite transform to correct for head-motion and susceptibility distortions. These resampled BOLD time-series will be referred to as preprocessed BOLD in original space, or just preprocessed BOLD. The BOLD time-series were resampled into standard space, generating a preprocessed BOLD run in MNI152NLin2009cAsym space. First, a reference volume and its skull-stripped version were generated using a custom methodology of fMRIPrep. Several confounding time-series were calculated based on the preprocessed BOLD: framewise displacement (FD), DVARS and three region-wise global signals. FD was computed using two formulations following Power (absolute sum of relative motions, Power et al. (2014)) and Jenkinson (relative root mean square displacement between affines, Jenkinson et al. (2002)). FD and DVARS are calculated for each functional run, both using their implementations in Nipype (following the definitions by Power et al. 2014). The three global signals are extracted within the CSF, the WM, and the whole-brain masks. Additionally, a set of physiological regressors were extracted to allow for component-based noise correction (CompCor, Behzadi et al. 2007). Principal components are estimated after high-pass filtering the preprocessed BOLD time-series (using a discrete cosine filter with 128s cut-off) for the two CompCor variants: temporal (tCompCor) and anatomical (aCompCor). tCompCor components are then calculated from the top 2% variable voxels within the brain mask. For aCompCor, three probabilistic masks (CSF, WM and combined CSF+WM) are generated in anatomical space. The implementation differs from that of Behzadi et al. in that instead of eroding the masks by 2 pixels on BOLD space, the aCompCor masks are subtracted a mask of pixels that likely contain a volume fraction of GM. This mask is obtained by dilating a GM mask extracted from the FreeSurfer's aseg segmentation, and it ensures components are not extracted from voxels containing a minimal fraction of GM. Finally, these masks are resampled into BOLD space and binarized by thresholding at 0.99 (as in the original implementation). Components are also calculated separately within the WM and CSF masks. For each CompCor decomposition, the k components with the largest singular values are retained, such that the retained components' time series are sufficient to explain 50 percent of variance across the nuisance mask (CSF, WM, combined, or temporal). The remaining components are dropped from consideration. The head-motion estimates calculated in the correction step were also placed within the corresponding confounds file. The confound time series derived from head motion estimates and global signals were expanded with the inclusion of temporal derivatives and quadratic terms for each (Satterthwaite et al. 2013). Frames that exceeded a threshold of 0.5 mm FD or 1.5 standardised DVARS were annotated as motion outliers. All resamplings can be performed with a single interpolation step by composing all the pertinent transformations (i.e. head-motion transform matrices, susceptibility distortion correction when available, and co-registrations to anatomical and output spaces). Gridded (volumetric) resamplings were performed using antsApplyTransforms (ANTs), configured with Lanczos interpolation to minimize the smoothing effects of other kernels (Lanczos 1964). Non-gridded (surface) resamplings were performed using mri_vol2surf (FreeSurfer). First, a reference volume and its skull-stripped version were generated using a custom methodology of fMRIPrep. A B0-nonuniformity map (or fieldmap) was estimated based on two (or more) echo-planar imaging (EPI) references with opposing phase-encoding directions, with 3dQwarp Cox and Hyde (1997) (AFNI 20160207). Based on the estimated susceptibility distortion, a corrected EPI (echo-planar imaging) reference was calculated for a more accurate co-registration with the anatomical reference. The BOLD reference was then co-registered to the T1w reference using bbregister (FreeSurfer) which implements boundary-based registration (Greve and Fischl 2009). Co-registration was configured with six degrees of freedom. Head-motion parameters with respect to the BOLD reference (transformation matrices, and six corresponding rotation and translation parameters) are estimated before any spatiotemporal filtering using mcflirt (FSL 5.0.9, Jenkinson et al. 2002). BOLD runs were slice-time corrected to 0.446s (0.5 of slice acquisition range 0s-0.892s) using 3dTshift from AFNI 20160207 (Cox and Hyde 1997, RRID:SCR_005927). The BOLD time-series (including slice-timing correction when applied) were resampled onto their original, native space by applying a single, composite transform to correct for head-motion and susceptibility distortions. These resampled BOLD time-series will be referred to as preprocessed BOLD in original space, or just preprocessed BOLD. The BOLD time-series were resampled into standard space, generating a preprocessed BOLD run in MNI152NLin2009cAsym space. First, a reference volume and its skull-stripped version were generated using a custom methodology of fMRIPrep. Several confounding time-series were calculated based on the preprocessed BOLD: framewise displacement (FD), DVARS and three region-wise global signals. FD was computed using two formulations following Power (absolute sum of relative motions, Power et al. (2014)) and Jenkinson (relative root mean square displacement between affines, Jenkinson et al. (2002)). FD and DVARS are calculated for each functional run, both using their implementations in Nipype (following the definitions by Power et al. 2014). The three global signals are extracted within the CSF, the WM, and the whole-brain masks. Additionally, a set of physiological regressors were extracted to allow for component-based noise correction (CompCor, Behzadi et al. 2007). Principal components are estimated after high-pass filtering the preprocessed BOLD time-series (using a discrete cosine filter with 128s cut-off) for the two CompCor variants: temporal (tCompCor) and anatomical (aCompCor). tCompCor components are then calculated from the top 2% variable voxels within the brain mask. For aCompCor, three probabilistic masks (CSF, WM and combined CSF+WM) are generated in anatomical space. The implementation differs from that of Behzadi et al. in that instead of eroding the masks by 2 pixels on BOLD space, the aCompCor masks are subtracted a mask of pixels that likely contain a volume fraction of GM. This mask is obtained by dilating a GM mask extracted from the FreeSurfer's aseg segmentation, and it ensures components are not extracted from voxels containing a minimal fraction of GM. Finally, these masks are resampled into BOLD space and binarized by thresholding at 0.99 (as in the original implementation). Components are also calculated separately within the WM and CSF masks. For each CompCor decomposition, the k components with the largest singular values are retained, such that the retained components' time series are sufficient to explain 50 percent of variance across the nuisance mask (CSF, WM, combined, or temporal). The remaining components are dropped from consideration. The head-motion estimates calculated in the correction step were also placed within the corresponding confounds file. The confound time series derived from head motion estimates and global signals were expanded with the inclusion of temporal derivatives and quadratic terms for each (Satterthwaite et al. 2013). Frames that exceeded a threshold of 0.5 mm FD or 1.5 standardised DVARS were annotated as motion outliers. All resamplings can be performed with a single interpolation step by composing all the pertinent transformations (i.e. head-motion transform matrices, susceptibility distortion correction when available, and co-registrations to anatomical and output spaces). Gridded (volumetric) resamplings were performed using antsApplyTransforms (ANTs), configured with Lanczos interpolation to minimize the smoothing effects of other kernels (Lanczos 1964). Non-gridded (surface) resamplings were performed using mri_vol2surf (FreeSurfer).

Many internal operations of fMRIPrep use Nilearn 0.6.2 (Abraham et al. 2014, RRID:SCR_001362), mostly within the functional processing workflow. For more details of the pipeline, see the section corresponding to workflows in fMRIPrep's documentation.

#### 7.2. Masks for exploring acoustic feature importance using regression analysis

##### 7.2.1. Valence network

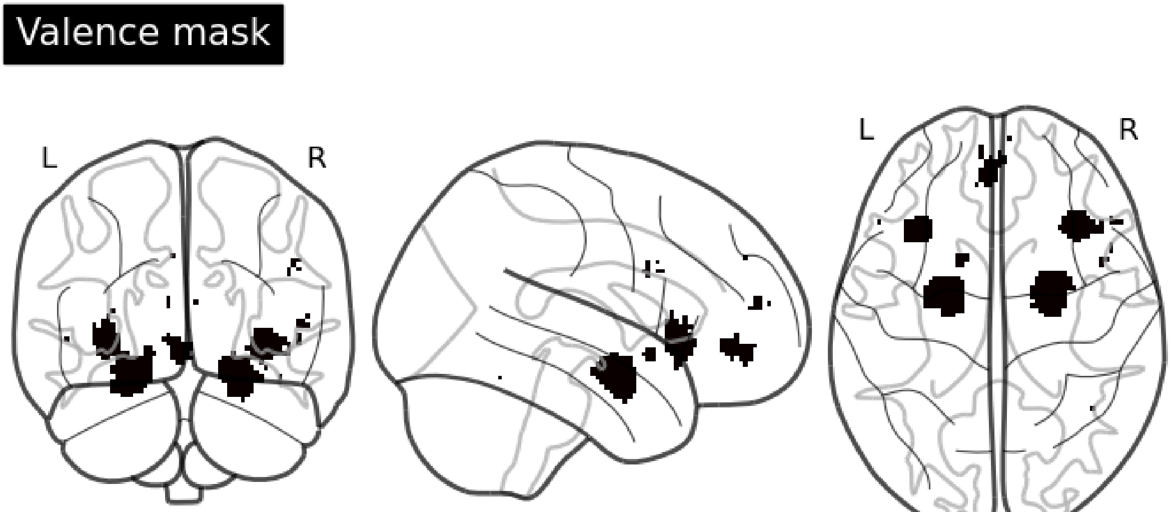


Figure S1 - Display of the results for the valence mask used in regression analysis.

Table S1 - Clusters identified in the map of Figure S1.

| Cluster ID | X | Y | Z | Peak Stat | Cluster Size (mm3) | structure |
|---|---|---|---|---|---|---|
| 1 | -32.5 | 21.5 | -2.5 | 1 | 1144 | Insula_L (aal) |
| 2 | -22.5 | -4.5 | -16.5 | 1 | 2832 | Amygdala_L (aal) |
| 3 | -14.5 | 9.5 | -8.5 | 1 | 112 | Putamen_L (aal) |
| 4 | -2.5 | 47.5 | -6.5 | 1 | 544 | Frontal_Med_Orb_L (aal) |
| 5 | -6.5 | 55.5 | 13.5 | 1 | 56 | Frontal_Sup_Medial_L (aal) |
| 6 | 23.5 | -4.5 | -18.5 | 1 | 2848 | Amygdala_R (aal) |
| 7 | 35.5 | 23.5 | -4.5 | 1 | 1336 | Insula_R (aal) |
| 8 | 51.5 | 25.5 | 3.5 | 1 | 48 | Frontal_Inf_Tri_R (aal) |

#### 7.2.2. Arousal mask

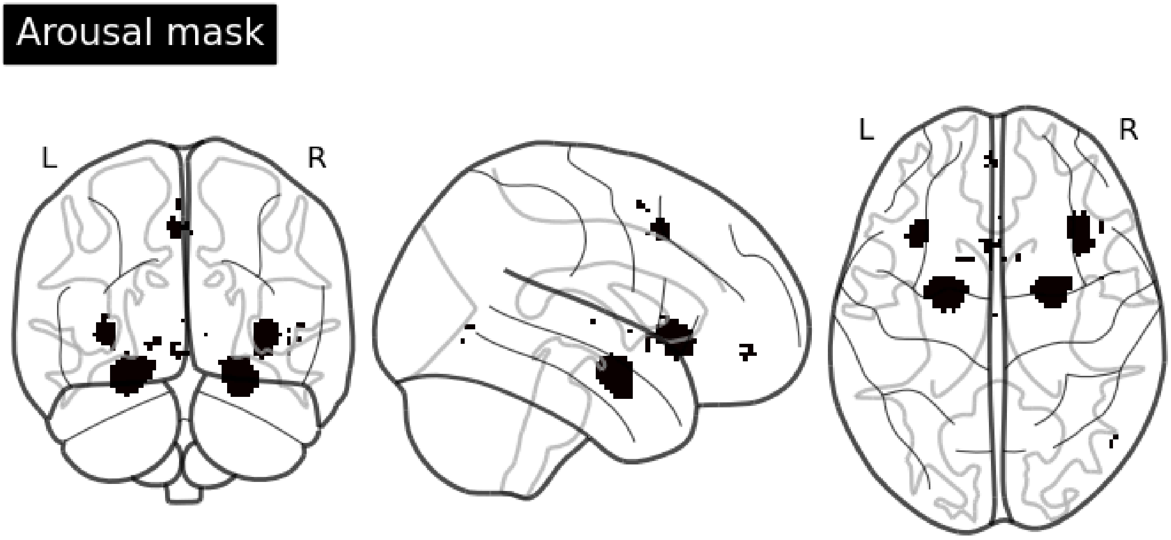


Figure S2 - Display of the results for the arousal mask used in regression analysis.

Table S2 - Clusters identified in the map of Figure S2.

| Cluster ID | X | Y | Z | Peak Stat | Cluster Size (mm3) | structure | |
|---|---|---|---|---|---|---|---|
| 0 | 1 | -32.5 | 21.5 | -0.5 | 1 | 664 | Insula_L (aal) |
| 1 | 2 | -22.5 | -4.5 | -16.5 | 1 | 2408 | Amygdala_L (aal) |
| 2 | 3 | -12.5 | 9.5 | -4.5 | 1 | 56 | Putamen_L (aal) |
| 3 | 4 | -4.5 | 15.5 | 45.5 | 1 | 240 | Supp_Motor_Area_L (aal) |
| 4 | 5 | -4.5 | 53.5 | -6.5 | 1 | 40 | Frontal_Med_Orb_L (aal) |
| 5 | 6 | 23.5 | -2.5 | -18.5 | 1 | 2056 | Amygdala_R (aal) |
| 6 | 7 | 35.5 | 21.5 | -2.5 | 1 | 1440 | Insula_R (aal) |
| 7 | 8 | 45.5 | 23.5 | -4.5 | 1 | 40 | Frontal_Inf_Orb_2_R (aal) |

##### 7.2.3. Primary auditory mask

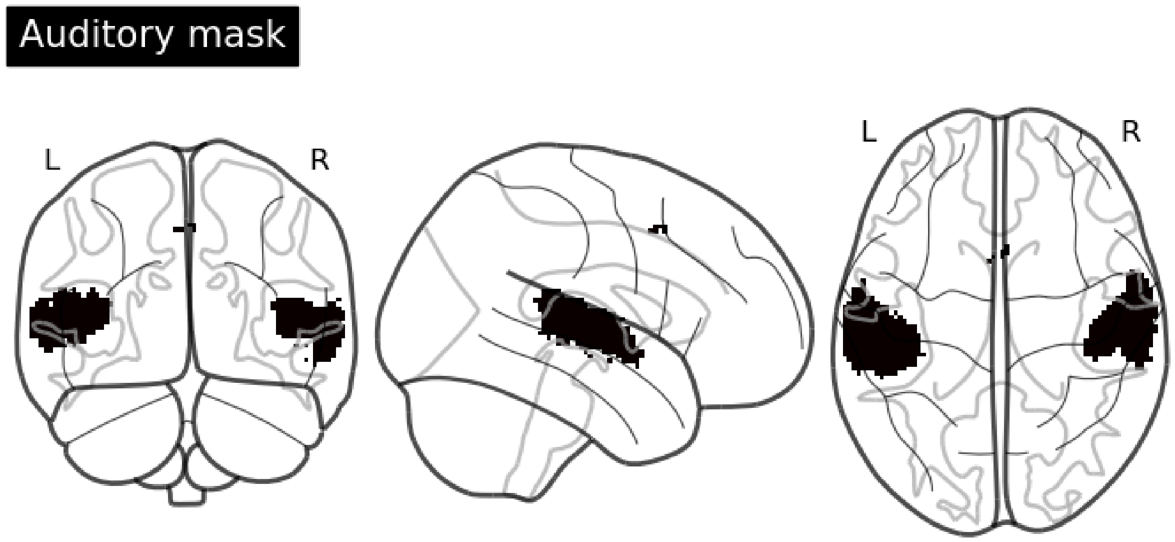


Figure S3 - Display of the results for the auditory mask used in regression analysis.

Table S3 - Clusters identified in the map of Figure S3.

| Cluster ID | X | Y | Z | Peak Stat | Cluster Size (mm3) | structure |
|---|---|---|---|---|---|---|
| 1 | -52.5 | -22.5 | 7.5 | 1 | 7600 | Temporal_Sup_L (aal) |
| 2 | -62.5 | -28.5 | 1.5 | 1 | 688 | Temporal_Mid_L (aal) |
| 3 | 3.5 | 13.5 | 47.5 | 1 | 40 | Supp_Motor_Area_R (aal) |
| 4 | 55.5 | -16.5 | 3.5 | 1 | 6592 | Temporal_Sup_R (aal) |

#### 7.2.4. Reward mask

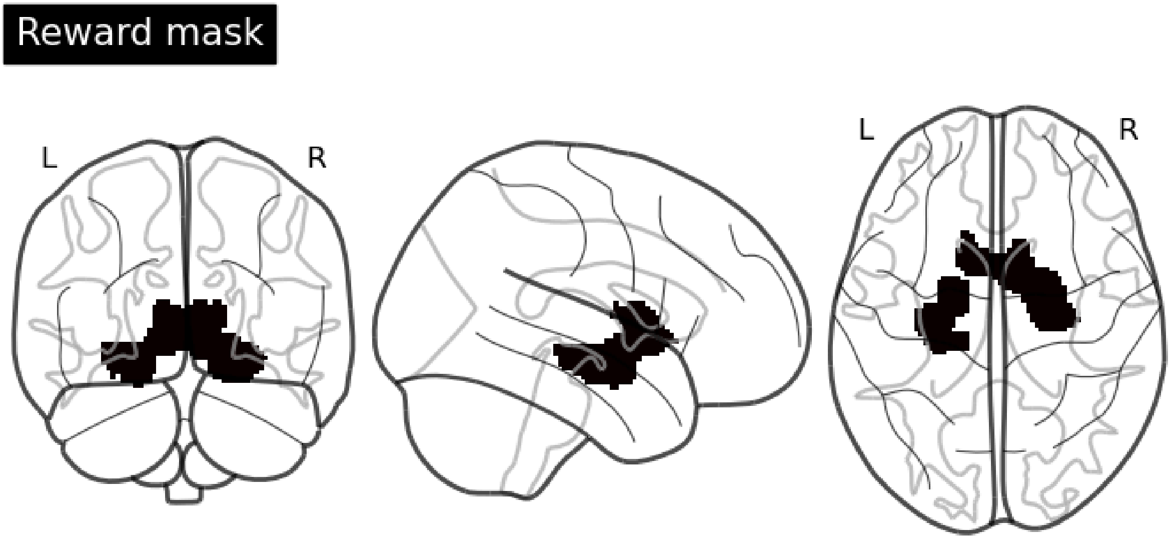


Figure S4 - Display of the results for the reward mask used in regression analysis.

Table S4 - Clusters identified in the map of Figure S4.

#### 7.3. MVPA results

##### 7.3.1. Classification per quadrant

Table S5 - Classification accuracies per quadrant and participant.

| Participant | Q1-acc | Q2-acc | Q3-acc | Q4-acc |
|---|---|---|---|---|
| 1 | 0.635417 | 0.635417 | 0.750000 | 0.666667 |
| 2 | 0.552083 | 0.729167 | 0.729167 | 0.687500 |
| 3 | 0.562500 | 0.729167 | 0.562500 | 0.708333 |
| 4 | 0.541667 | 0.666667 | 0.718750 | 0.687500 |
| 7 | 0.645833 | 0.750000 | 0.781250 | 0.781250 |
| 8 | 0.531250 | 0.645833 | 0.843750 | 0.843750 |
| 9 | 0.656250 | 0.718750 | 0.729167 | 0.697917 |
| 10 | 0.572917 | 0.812500 | 0.770833 | 0.666667 |
| 11 | 0.583333 | 0.906250 | 0.750000 | 0.666667 |
| 12 | 0.750000 | 0.666667 | 0.687500 | 0.583333 |
| 13 | 0.708333 | 0.708333 | 0.635417 | 0.770833 |
| 14 | 0.510417 | 0.854167 | 0.760417 | 0.708333 |
| 15 | 0.395833 | 0.937500 | 0.750000 | 0.572917 |
| 16 | 0.479167 | 0.875000 | 0.937500 | 0.604167 |
| 17 | 0.468750 | 0.656250 | 0.729167 | 0.812500 |
| 19 | 0.697917 | 0.750000 | 0.854167 | 0.625000 |

#### 7.3.2. Maps with most important voxels

### 7.3.2.1. Q1

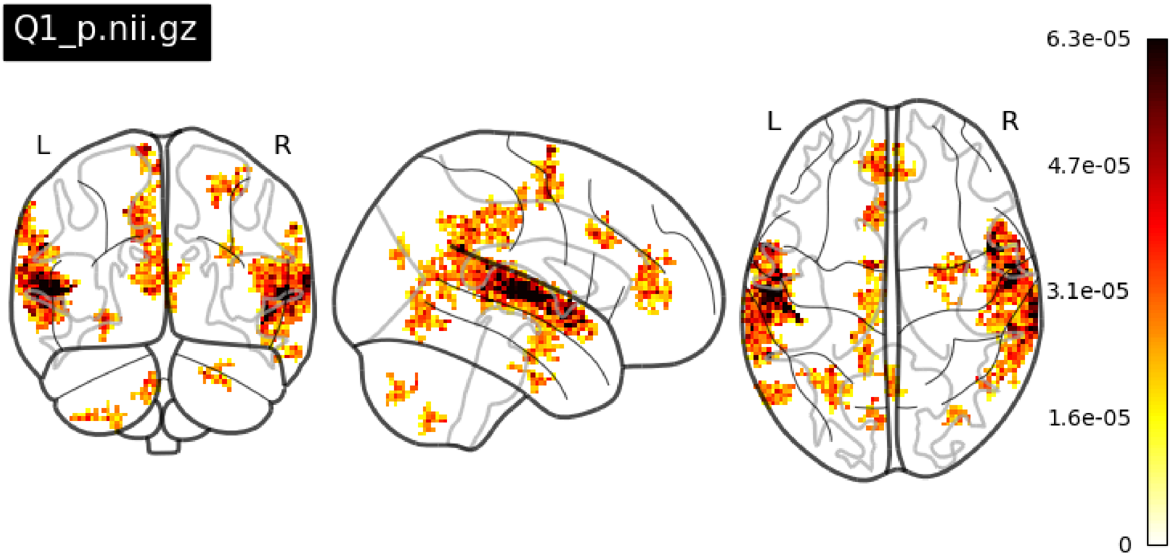


Figure S5 - Glass brain plot with the most discriminant voxels for quadrant Q1.

### 7.3.2.2. Q2

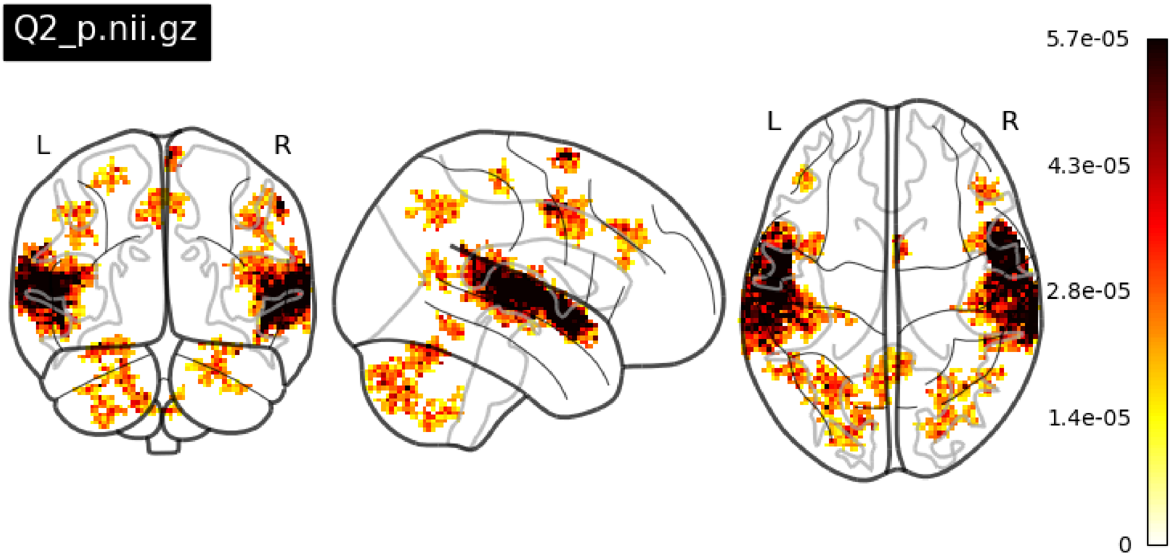


Figure S6 - Glass brain plot with the most discriminant voxels for quadrant Q2.

### 7.3.2.3. Q3

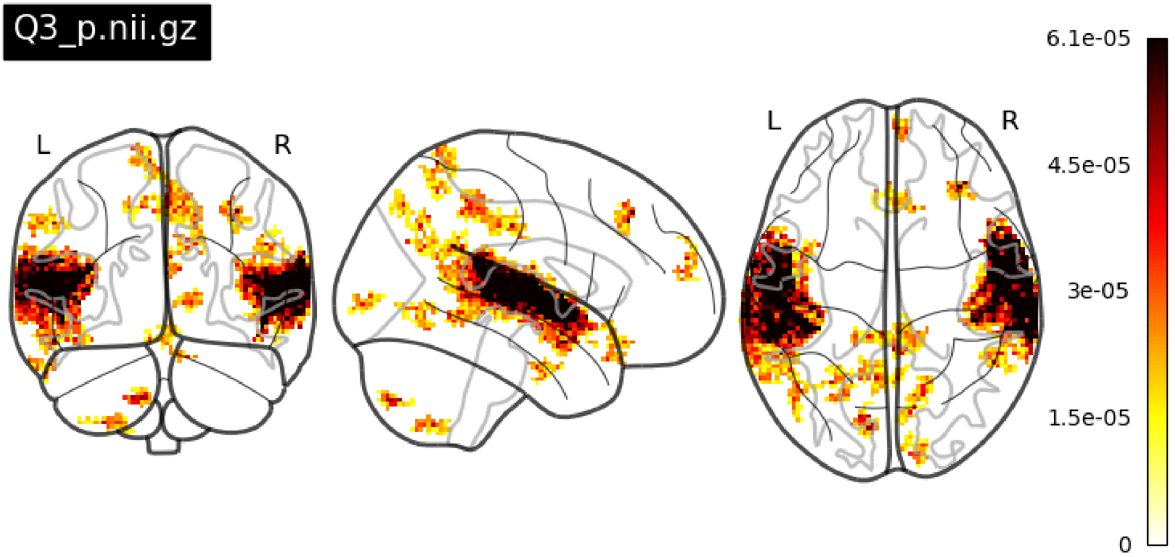


Figure S7 - Glass brain plot with the most discriminant voxels for quadrant Q2.

### 7.3.2.4. Q4

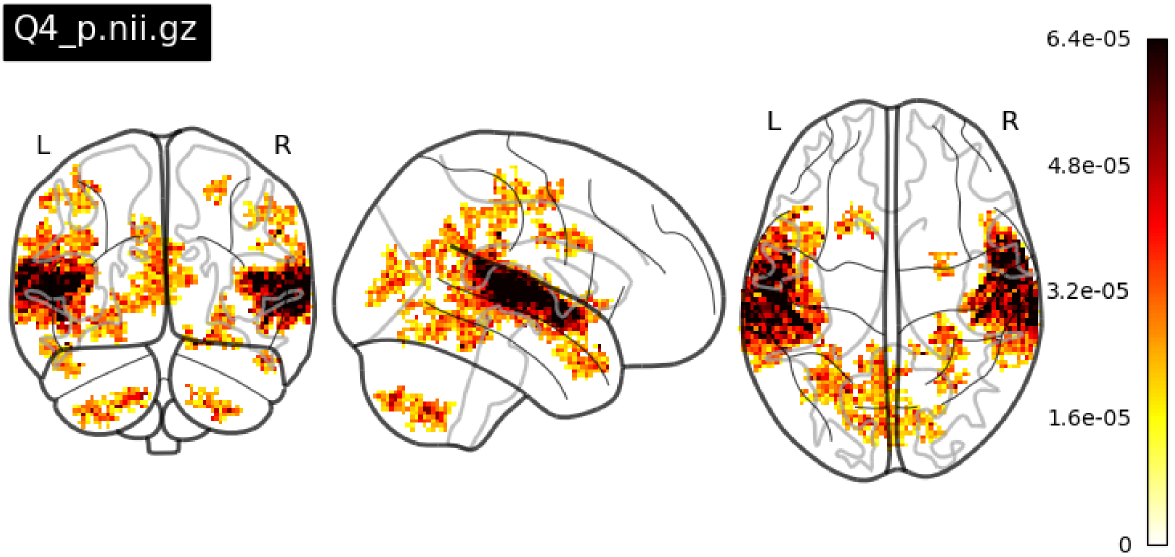


Figure S8 - Glass brain plot with the most discriminant voxels for quadrant Q4.

### 7.4. Regression analysis of acoustical features

#### 7.4.1. Valence meta-analysis mask

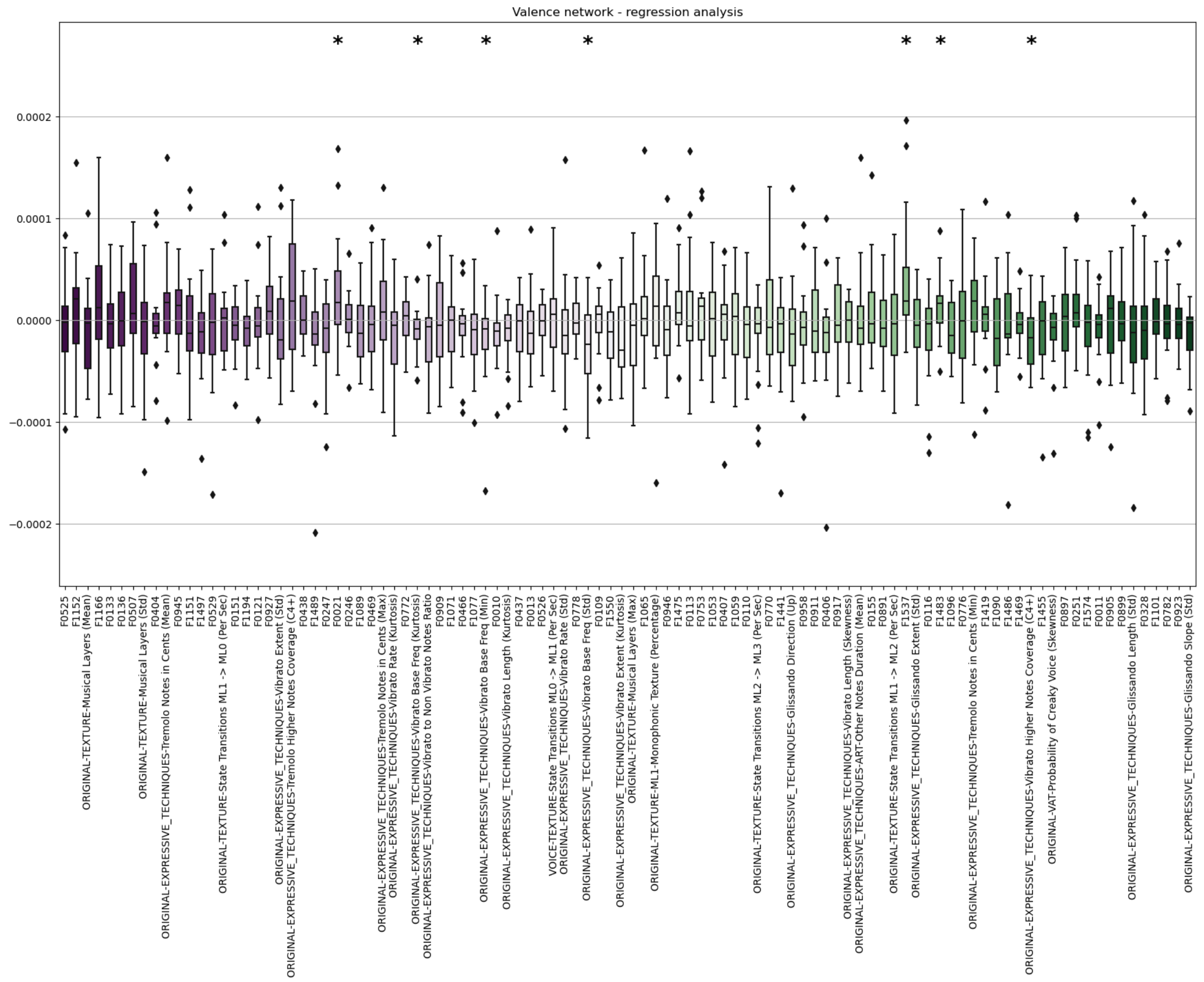


Figure S9 - Regression coefficients for the 100 features considering the valence mask.

Table S6 - Identification of the features with significant coefficients for the valence mask.

| name | description | p-value |
|---|---|---|
| F0021 | Events Density | 0.04 |
| ORIGINAL-EXPRESSIVE_TECHNIQUES-Vibrato Base Freq (Kurtosis) | NA | 0.026 |
| ORIGINAL-EXPRESSIVE_TECHNIQUES-Vibrato Base Freq (Min) | NA | 0.04 |
| ORIGINAL-EXPRESSIVE_TECHNIQUES-Vibrato Base Freq (Std) | NA | 0.036 |
| F1537 | Pitch (Terhardt) - Chord Change Likelihood (median) | 0.02 |
| F1483 | Loudness (MG & B PsySound2) - Tonal Dissonance (S) (skewness) | 0.036 |
| ORIGINAL-EXPRESSIVE_TECHNIQUES-Vibrato Higher Notes Coverage (C4+) | NA | 0.036 |

#### 7.4.2. Arousal meta-analysis mask

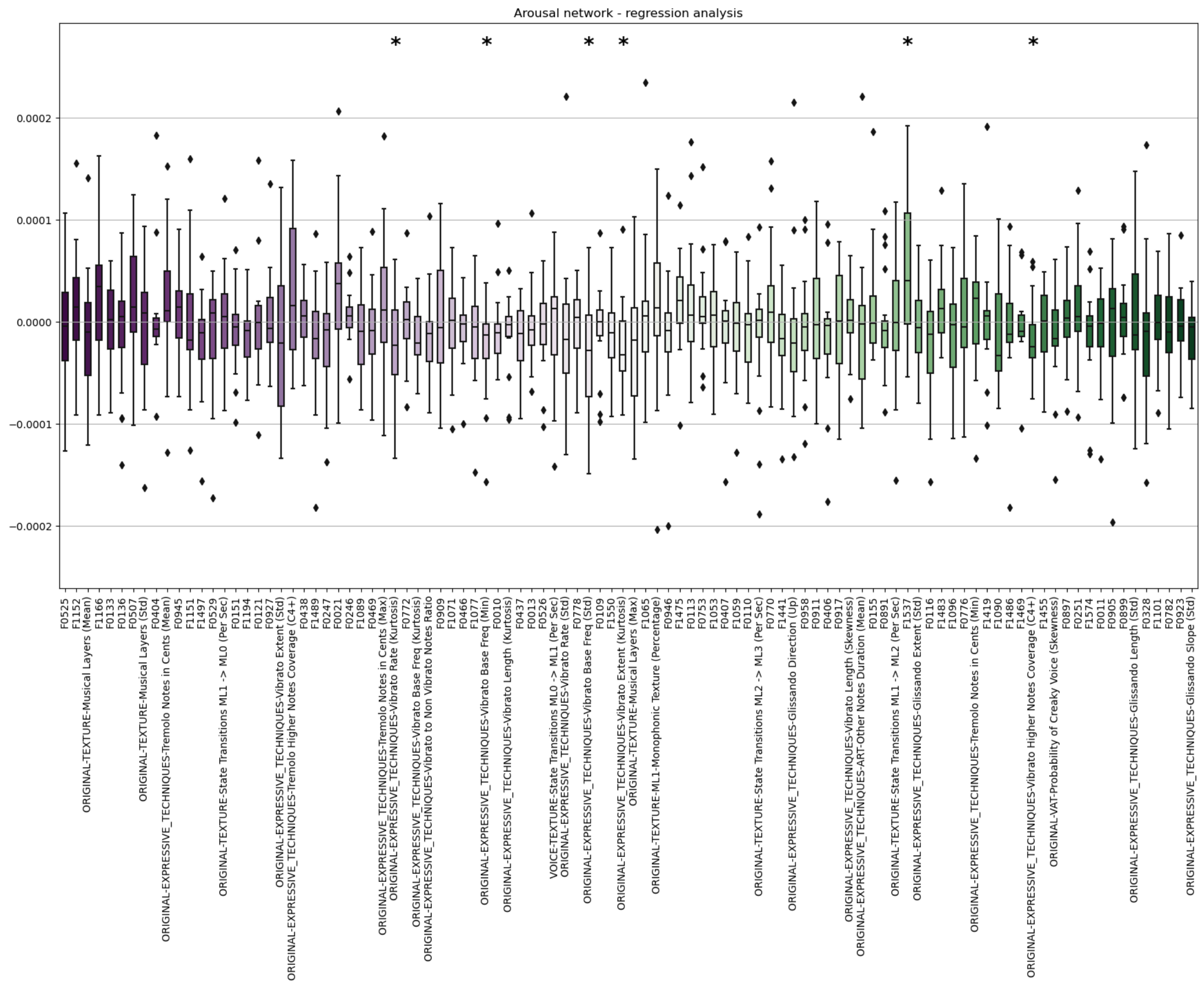


Figure S10 - Regression coefficients for the 100 features considering the arousal mask.

Table S7 - Identification of the features with significant coefficients for the arousal mask.

| name | description | p-value |
|---|---|---|
| ORIGINAL-EXPRESSIVE_TECHNIQUES-Vibrato Rate (Kurtosis) | NA | 0.032 |
| ORIGINAL-EXPRESSIVE_TECHNIQUES-Vibrato Base Freq (Min) | NA | 0.011 |
| ORIGINAL-EXPRESSIVE_TECHNIQUES-Vibrato Base Freq (Std) | NA | 0.049 |
| ORIGINAL-EXPRESSIVE_TECHNIQUES-Vibrato Extent (Kurtosis) | NA | 0.014 |
| F1537 | Pitch (Terhardt) - Chord Change Likelihood (median) | 0.009 |
| ORIGINAL-EXPRESSIVE_TECHNIQUES-Vibrato Higher Notes Coverage (C4+) | NA | 0.04 |

##### 7.4.3. Primary auditory meta-analysis mask

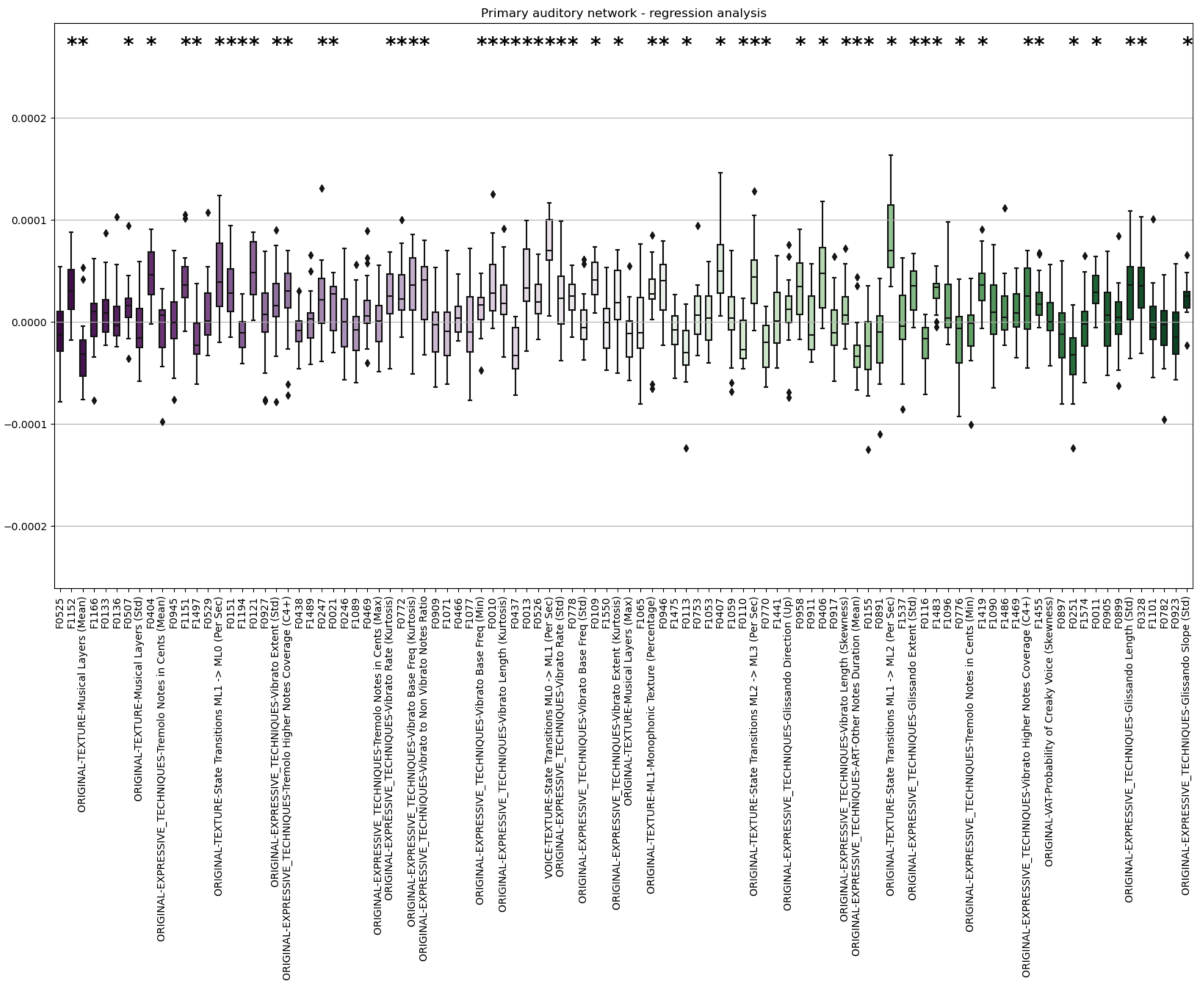


Figure S11 - Regression coefficients for the 100 features considering the auditory mask.

Table S8- Identification of the features with significant coefficients for the auditory mask.

| name | description | p-value |
|---|---|---|
| F1152 | FFT Spectrum - Average Power Spectrum (median) | 0.001 |
| ORIGINAL-TEXTURE-Musical Layers (Mean) | NA | 0.003 |
| F0507 | Rolloff (mean) | 0.006 |
| F0404 | Rolloff (MeanA/StdM) | 0.0 |
| F1151 | FFT Spectrum - Average Power Spectrum (mean) | 0.0 |
| F1497 | Loudness (MG & B PsySound2) - Spectral Dissonance (S) (skewness) | 0.011 |
| ORIGINAL-TEXTURE-State Transitions ML1 -> ML0 (Per Sec) | NA | 0.0 |
| F0151 | Spectral Entropy (std) | 0.0 |
| F1194 | FFT Spectrum - Skewness (median) | 0.023 |
| F0121 | Spectral Centroid (std) | 0.0 |
| ORIGINAL-EXPRESSIVE_TECHNIQUES-Vibrato Extent (Std) | NA | 0.026 |
| ORIGINAL-EXPRESSIVE_TECHNIQUES-Tremolo Higher Notes Coverage (C4+) | NA | 0.045 |
| F0247 | Inharmonicity (std) | 0.007 |
| F0021 | Events Density | 0.012 |
| ORIGINAL-EXPRESSIVE_TECHNIQUES-Vibrato Rate (Kurtosis) | NA | 0.003 |
| F0772 | LSP_4 (std) | 0.0 |
| ORIGINAL-EXPRESSIVE_TECHNIQUES-Vibrato Base Freq (Kurtosis) | NA | 0.0 |
| ORIGINAL-EXPRESSIVE_TECHNIQUES-Vibrato to Non Vibrato Notes Ratio | NA | 0.001 |
| ORIGINAL-EXPRESSIVE_TECHNIQUES-Vibrato Base Freq (Min) | NA | 0.011 |
| F0010 | Fluctuation (std) | 0.0 |
| ORIGINAL-EXPRESSIVE_TECHNIQUES-Vibrato Length (Kurtosis) | NA | 0.011 |
| F0437 | MFCC0 (StdA/MeanM) | 0.0 |

| | | |
|---|---|---|
| F0013 | Fluctuation (max) | 0.0 |
| F0526 | MFCC1 (std) | 0.0 |
| VOICE-TEXTURE-State Transitions ML0 -> ML1 (Per Sec) | NA | 0.0 |
| ORIGINAL-EXPRESSIVE_TECHNIQUES-Vibrato Rate (Std) | NA | 0.012 |
| F0778 | LSP_5 (std) | 0.001 |
| F0109 | High-frequency Energy (std) | 0.0 |
| ORIGINAL-EXPRESSIVE_TECHNIQUES-Vibrato Extent (Kurtosis) | NA | 0.011 |
| ORIGINAL-TEXTURE-ML1-Monophonic Texture (Percentage) | NA | 0.011 |
| F0946 | SFM_15 (std) | 0.001 |
| F0113 | High-frequency Energy (min) | 0.001 |
| F0407 | MFCC1 (MeanA/StdM) | 0.0 |
| F0110 | High-frequency Energy (skewness) | 0.004 |
| ORIGINAL-TEXTURE-State Transitions ML2 -> ML3 (Per Sec) | NA | 0.0 |
| F0770 | LSP_3 (min) | 0.001 |
| F0958 | SFM_17 (std) | 0.0 |
| F0406 | MFCC0 (MeanA/StdM) | 0.0 |
| ORIGINAL-EXPRESSIVE_TECHNIQUES-Vibrato Length (Skewness) | NA | 0.02 |
| ORIGINAL-EXPRESSIVE_TECHNIQUES-ART-Other Notes Duration (Mean) | NA | 0.003 |
| F0155 | Spectral Entropy (min) | 0.005 |
| ORIGINAL-TEXTURE-State Transitions ML1 -> ML2 (Per Sec) | NA | 0.0 |
| ORIGINAL-EXPRESSIVE_TECHNIQUES-Glissando Extent (Std) | NA | 0.0 |
| F0116 | Spectral Flux (skewness) | 0.0 |
| F1483 | Loudness (MG & B PsySound2) - Tonal Dissonance (S) (skewness) | 0.0 |
| F0776 | LSP_4 (min) | 0.045 |
| F1419 | Dynamic Loudness (C & F) - Sharpness (std) | 0.0 |

| | | |
|---|---|---|
| ORIGINAL-EXPRESSIVE_TECHNIQUES-Vibrato Higher Notes Coverage (C4+) | NA | 0.029 |
| F1455 | Loudness (MG & B PsySound2) - SharpnessA (skewness) | 0.0 |
| F0251 | Inharmonicity (min) | 0.0 |
| F0011 | Fluctuation (skewness) | 0.0 |
| ORIGINAL-EXPRESSIVE_TECHNIQUES-Glissando Length (Std) | NA | 0.003 |
| F0328 | Tonal Centroid 3 (std) | 0.0 |
| ORIGINAL-EXPRESSIVE_TECHNIQUES-Glissando Slope (Std) | NA | 0.0 |

#### 7.4.4. Reward meta-analysis mask

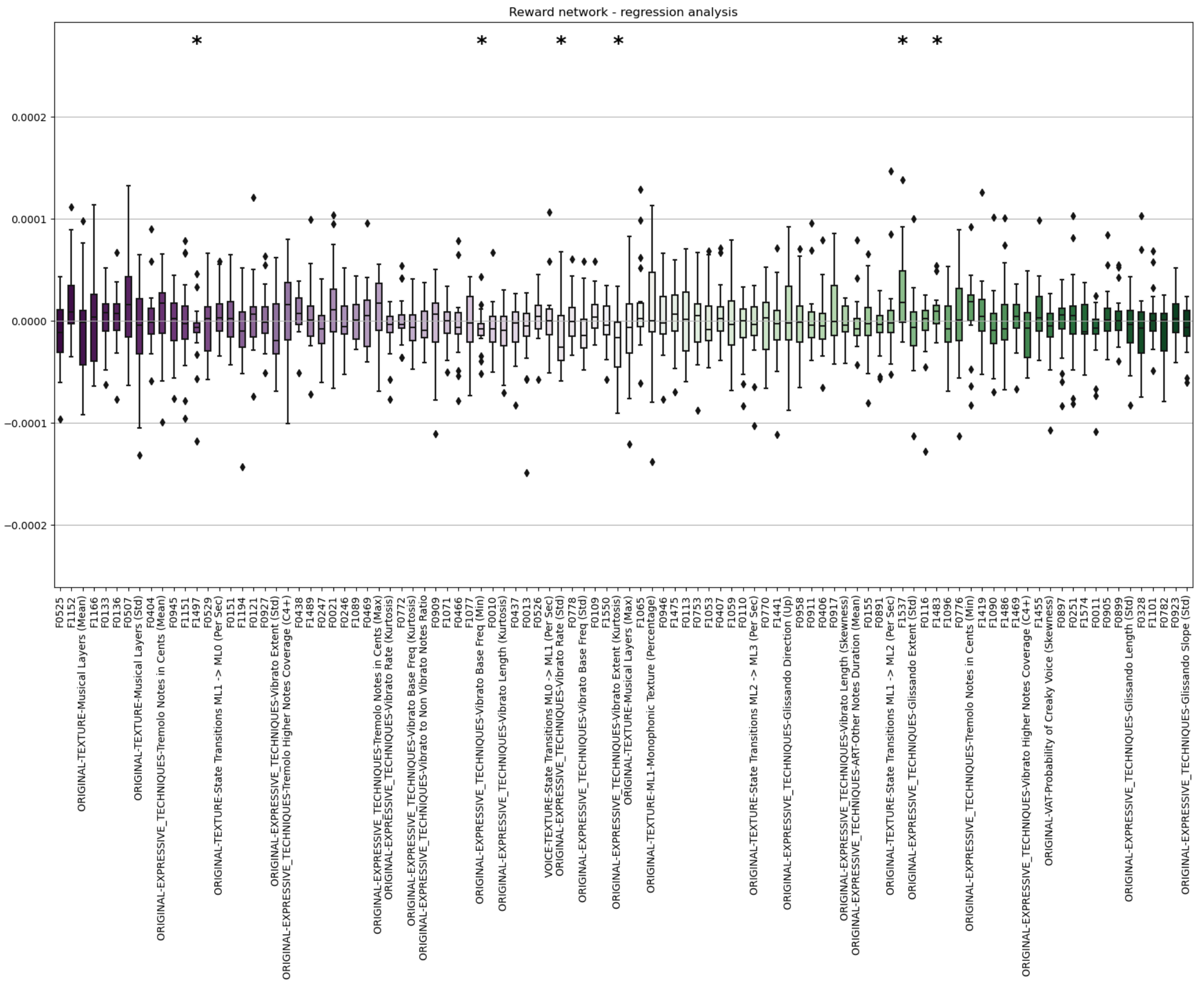


Figure S12 - Regression coefficients for the 100 features considering the reward mask.

Table S9 - Identification of the features with significant coefficients for the reward mask.

| name | description | p-value |
|---|---|---|
| F1497 | Loudness (MG & B PsySound2) - Spectral Dissona... | 0.032 |
| ORIGINAL-EXPRESSIVE_TECHNIQUES -Vibrato Base Fr... | NA | 0.036 |
| ORIGINAL-EXPRESSIVE_TECHNIQUES -Vibrato Rate (Std) | NA | 0.040 |
| ORIGINAL-EXPRESSIVE_TECHNIQUES -Vibrato Extent ... | NA | 0.018 |
| F1537 | Pitch (Terhardt) - Chord Change Likelihood (me... | 0.005 |
| F1483 | Loudness (MG & B PsySound2) - Tonal Dissonance... | 0.045 |

#### 7.5. POMS

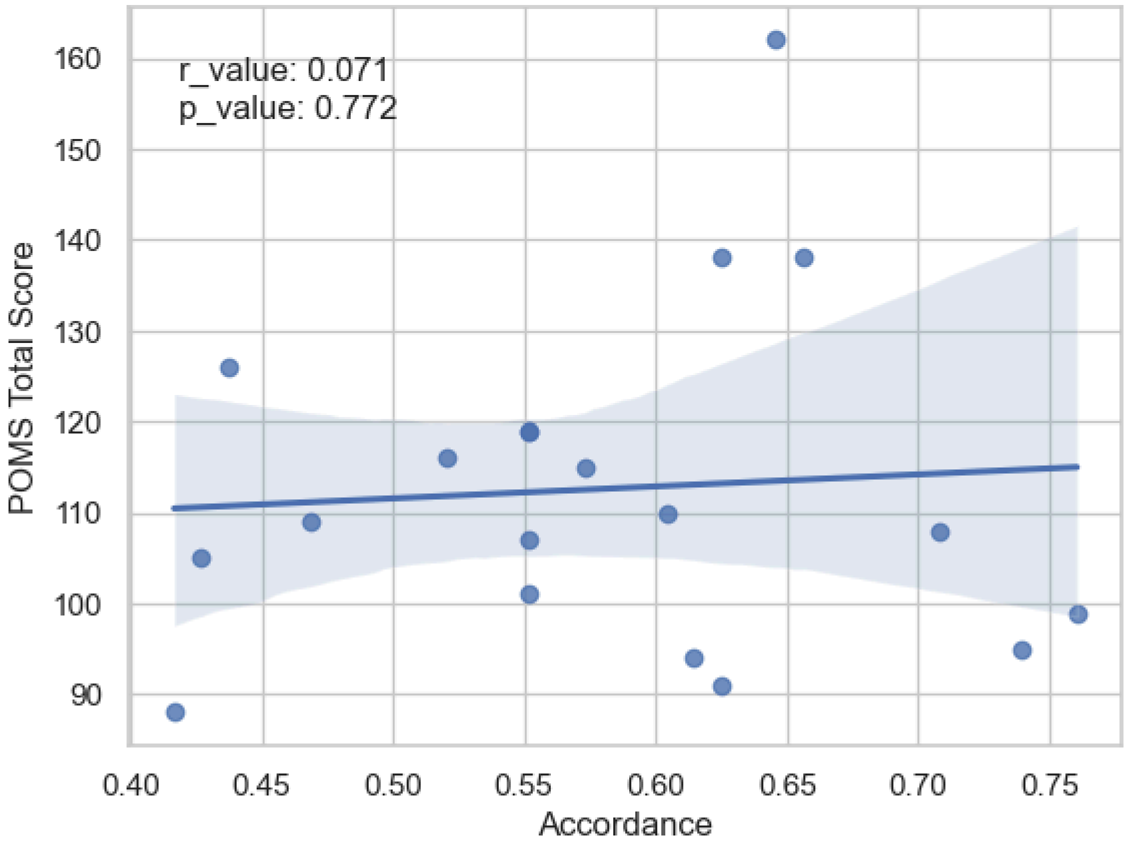


Figure S13 - Regression analysis between the accordance in the quadrant rating (database label - participant label) and the POMS total score before the session.

#### 7.6. Z-score comparison for arousal and valence

The test for Arousal groups the Q1 and Q2 labels as 'positive' and Q3 and Q4 as 'negative'. The test for valence groups the Q1 and Q4 labels as 'positive' and Q2 and Q3 as 'negative'. We compared the z-values between positive and negative labels for all clusters using a two-sided Mann-Whitney-Wilcoxon test with FDR correction (q=0.05).

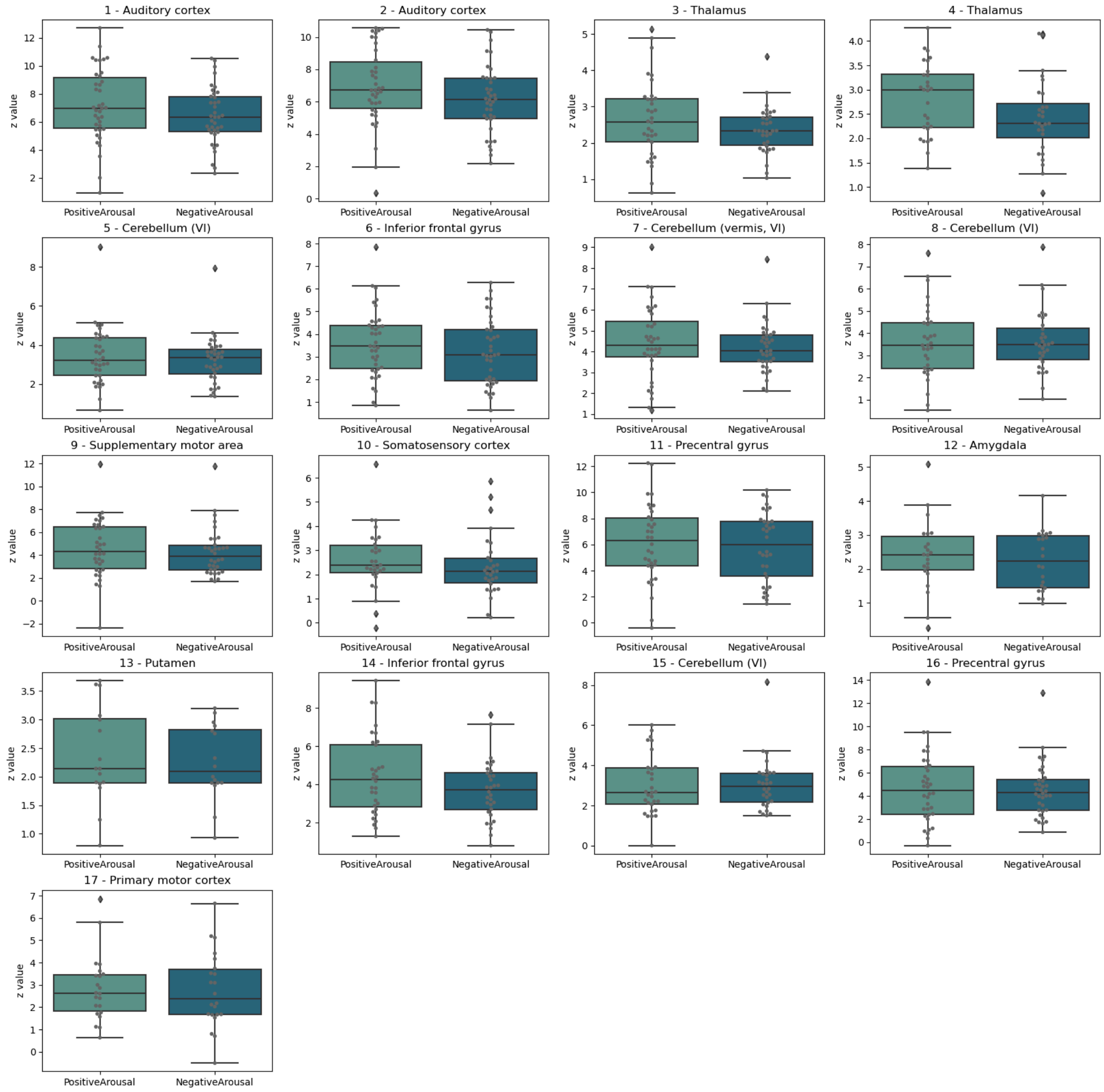


Figure S14 - Boxplots for the z values of each identified cluster (Table 1) divided per positive and negative arousal quadrants. Pairwise statistical comparisons between quadrants are displayed, performed using two-sided Mann-Whitney-Wilcoxon tests with FDR correction (q=0.05).

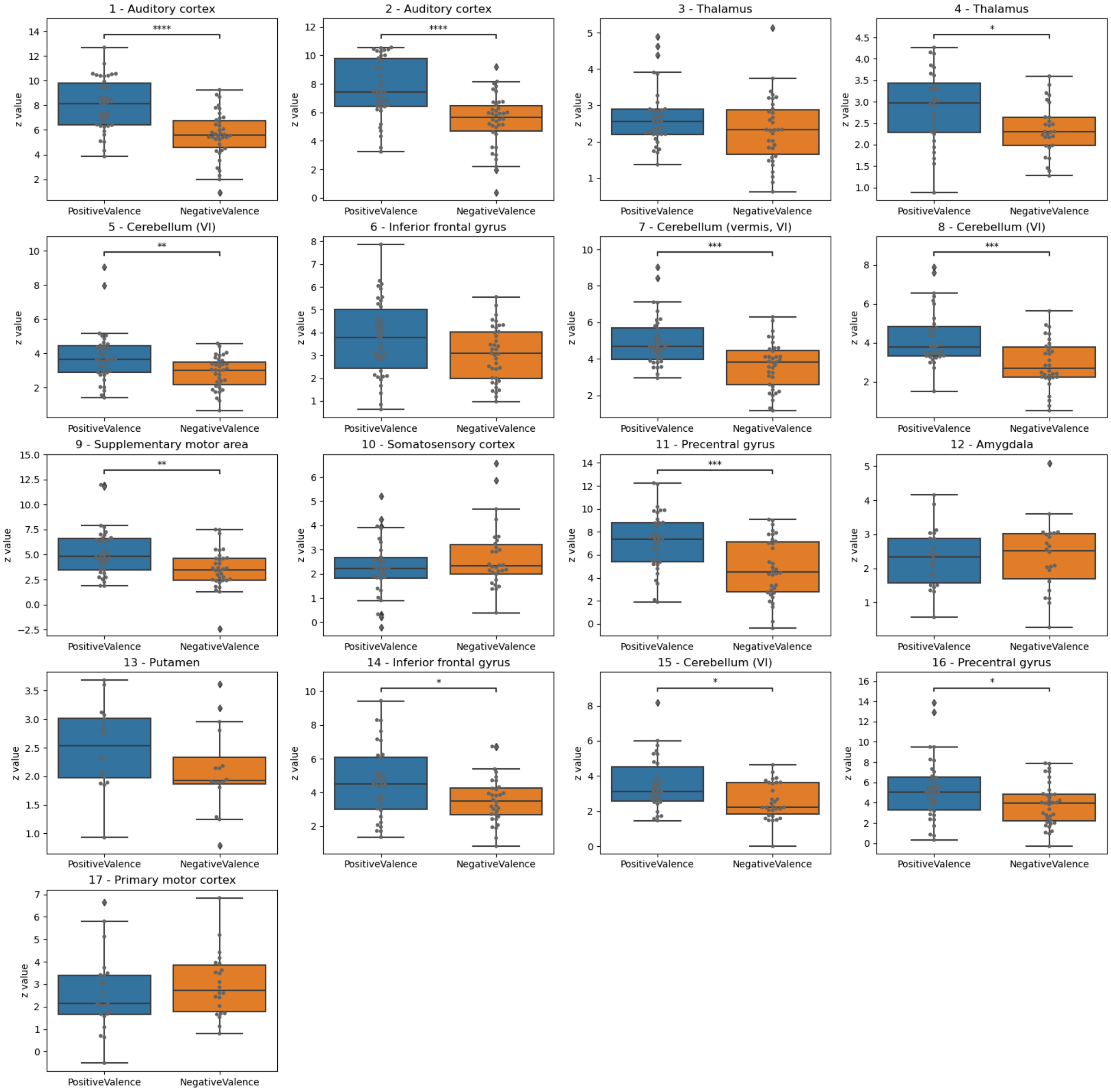


Figure S15 - Boxplots for the z values of each identified cluster (Table 1) divided per positive and negative valence quadrants. Pairwise statistical comparisons between quadrants are displayed, performed using two-sided Mann-Whitney-Wilcoxon tests with FDR correction (q=0.05).
